# Spatial transcriptomic analysis drives PET imaging of tight junction protein expression in pancreatic cancer theranostics

**DOI:** 10.1101/2024.01.07.574209

**Authors:** James Wang, Jai Woong Seo, Aris J. Kare, Martin Schneider, Spencer K. Tumbale, Bo Wu, Marina N. Raie, Mallesh Pandrala, Andrei Iagaru, Ryan L. Brunsing, Gregory W. Charville, Walter G. Park, Katherine W. Ferrara

## Abstract

We apply spatial transcriptomics and proteomics to select pancreatic cancer surface receptor targets for molecular imaging and theranostics using an approach that can be applied to many cancers. Selected cancer surfaceome epithelial markers were spatially correlated and provided specific cancer localization, whereas the spatial correlation between cancer markers and immune- cell or fibroblast markers was low. While molecular imaging of cancer-associated fibroblasts and integrins has been proposed for pancreatic cancer, our data point to the tight junction protein claudin-4 as a theranostic target. Claudin-4 expression increased ∼16 fold in cancer as compared with normal pancreas, and the tight junction localization conferred low background for imaging in normal tissue. We developed a peptide-based molecular imaging agent targeted to claudin-4 with accumulation to ∼25% injected activity per cc (IA/cc) in metastases and ∼18% IA/cc in tumors. Our work motivates a new approach for data-driven selection of molecular targets.

## Introduction

Molecular Imaging (MI) is a growing biomedical discipline that enables the non-invasive visualization, characterization and quantification of chemical and biological processes at the cellular and subcellular levels across time in living systems. Positron emission tomography (PET) uniquely quantifies biological function by employing sub-pharmacological amounts of molecularly-specific radiolabeled agents (radiopharmaceuticals). Their short radioactive half-lives facilitate repeated imaging, and the latest hardware innovations enable dynamic functional imaging of these biomarkers *in situ*. Molecular imaging can detect small cancerous lesions and provides an opportunity to assess therapeutic accumulation across a wide range of molecular therapies. Cell surface receptors are preferred for imaging and therapy due to the relative efficiency of delivery, where an ideal agent will safely accumulate within minutes while excess agent is rapidly cleared from the body, facilitating imaging and minimally-toxic therapies.

The selection of receptor targets for molecular imaging and therapies has lacked a data-driven basis for the comparative evaluation of target receptors. Here, we address this important goal by applying spatial transcriptomics and proteomics to select molecular imaging agents. We apply these methods in pancreatic ductal adenocarcinoma (PDAC), but the methods developed here are general and can be applied across a range of disease targets. Characterizing pancreatic nodules with imaging is significant as 30-50% of the adult population may harbor precancerous lesions ^1–4^, invasive cancer can be present within larger intraductal papillary mucinous neoplasms (IPMNs) and incidentally-discovered pancreatic intraepithelial neoplasm (PanIN) precancers can include similar mutational profiles to PDAC ^2^. Further, PDAC is the seventh leading cause of cancer-related deaths as of 2021 ^5^.

In order to improve patient outcomes, specific imaging methods to detect disease and guide therapy are needed. PET-CT imaging has been applied using ^18^F-FDG imaging of metabolism ^6–8^; however, ^18^F-FDG has low sensitivity and specificity in the diagnosis of PDAC with variable detection of metastatic lymph nodes and false-positive findings in inflammation ^9,10^. Further, new strategies are required to perform noninvasive tumor, lymph node, and metastasis evaluations ^9^. Alternative markers currently under study, including integrins and fibroblast markers are expressed by only a fraction of tumor cells (e.g. ITGB6, in 49% of PDAC cells^11^) and have substantial background in many cases ^11^. We explore the spatial distribution of such markers and compare with cancer epithelial alternatives here.

Spatial mRNA transcriptomics and proteomics are ideal for identifying theranostic targets due to the combination of spatial and biological insight. More recently, cancer cell surface proteins (SPs) have been detailed for multiple cancers with comprehensive identification and annotation of genes encoding SPs (GESPs) via pan-cancer analysis. We employ spatial transcriptomics to survey mRNA within a multi-centimeter region of interest in formalin-fixed, paraffin-embedded (FFPE) tissue from human surgical samples with a depth of ∼18,000 genes. We combine spatial transcriptomics with multiplexed spatial proteomics (codetection by indexing, CODEX) or immunohistochemistry (IHC) to characterize PDAC, precancer, and surrounding pancreas from FFPE human surgical samples.

Claudin-4 (CLDN4) is a particularly attractive target as it is a member of a family of tight junction transmembrane proteins with increased expression in PDAC, and we hypothesized that the protein would be exposed to a radioligand only in the presence of disease ^12–14^. CLDN4 expression is also enhanced in prostate ^15^, breast ^16^ and ovarian ^17^ cancers and as pancreatic precancers advance, the expression level of *CLDN4* increases (p=0.001) ^12^. Intense positive CLDN4 immunolabeling has been noted within virtually all primary (71/72 [99%]) and metastatic (49/49 [100%]) PDAC tissue samples and in 10 of 11 precancerous PanIN specimens ^18^. Although claudin 6 and 18.2 have been the basis for development of therapeutics for ovarian and esophageal/gastric cancers ^19^, respectively, CLDN4 represents an additional promising therapeutic target. Claudin-18.2 is expressed in only a fraction of PDAC cases, as compared with 90-100% for CLDN4 ^11,18^.

Our paper is organized as follows. In human surgical specimens, we first evaluate the spatial correlation of mRNA expression with cell surface markers known to be upregulated in PDAC and precancers. We further evaluate the pseudotime evolution of the precancers and cancer to assess the evolution of candidate receptors. Single-cell evaluation of protein expression in PDAC and precancers is then conducted, providing a view of epithelial markers, fibroblasts and immune cells with a higher spatial resolution. Finally, we apply the results from transcriptomics and proteomics to select CLDN4 as an imaging marker and assess the resulting imaging in multiple models of murine PDAC.

## Results

### Spatial sequencing of PDAC samples elucidates spatially-correlated cancer surfaceome markers

Our discovery cohort includes 40 human surgical samples of surrounding pancreas, IPMNs^20^ or PDAC^21^ acquired at Stanford or from available databases ^20,21^ (Supplementary Table S1, Fig. 1A) and single-annotated cells from references ^21–23^. All human data collection and analysis was approved by the Stanford Institutional Review Board. The spatial transcriptomics samples span precancers, naïve untreated PDAC and patients treated with chemotherapy or chemo-radiation or mixed treatment. We first integrated data from treatment-naïve PDAC samples and performed unsupervised clustering (Supplementary Fig. S1, Fig. 1A-B). Top genes in all spatial clusters are summarized in Supplementary Fig. S1B, whereas Fig. 1B-H focuses on the top surfaceome atlas markers in the cancer cluster. Cluster 1 in Fig. 1C was enriched in cells with an epithelial, EpCAM^+^ Duct, and gastric mature pit phenotype (Fig. 1C)^24^. In particular, Cluster 1 differentially expressed *S100P, MUC5AC, TFF1* and *CLDN4* as compared with other clusters and compared with surrounding normal pancreas (Fig. 1D). To quantify how top cancer surfaceome atlas markers change across clusters (Supplementary Table S2), we created a cancer surfaceome score. The cancer cell surfaceome expression levels differed among clusters and were upregulated in Cluster 1 (Fig. 1E). The correlation of a set of the cancer surfaceome epithelial markers with one another (*CLDN4, GRPC5A, S100P, TSPAN8, MUC1, TFF1*) was >0.6 averaged across all spatial locations in the naïve PDAC cohort (Fig. 1F). Merging scaled and normalized data across transcriptomic sets from Stanford and HTAN, the fraction of spots in Cluster 1 overexpressing *CLDN4, S100P, TFF1, CEACAM5, MUC5AC* and *FAP* was 89, 78, 57, 46, 60 and 27%, respectively. With integration of batch-corrected data from Stanford and HTAN^21^ (rather than merging the datasets across sites), the fraction of spots with *CLDN4* overexpression increased to 97.5%, and differences across key genes were consistent.

**Figure 1.**
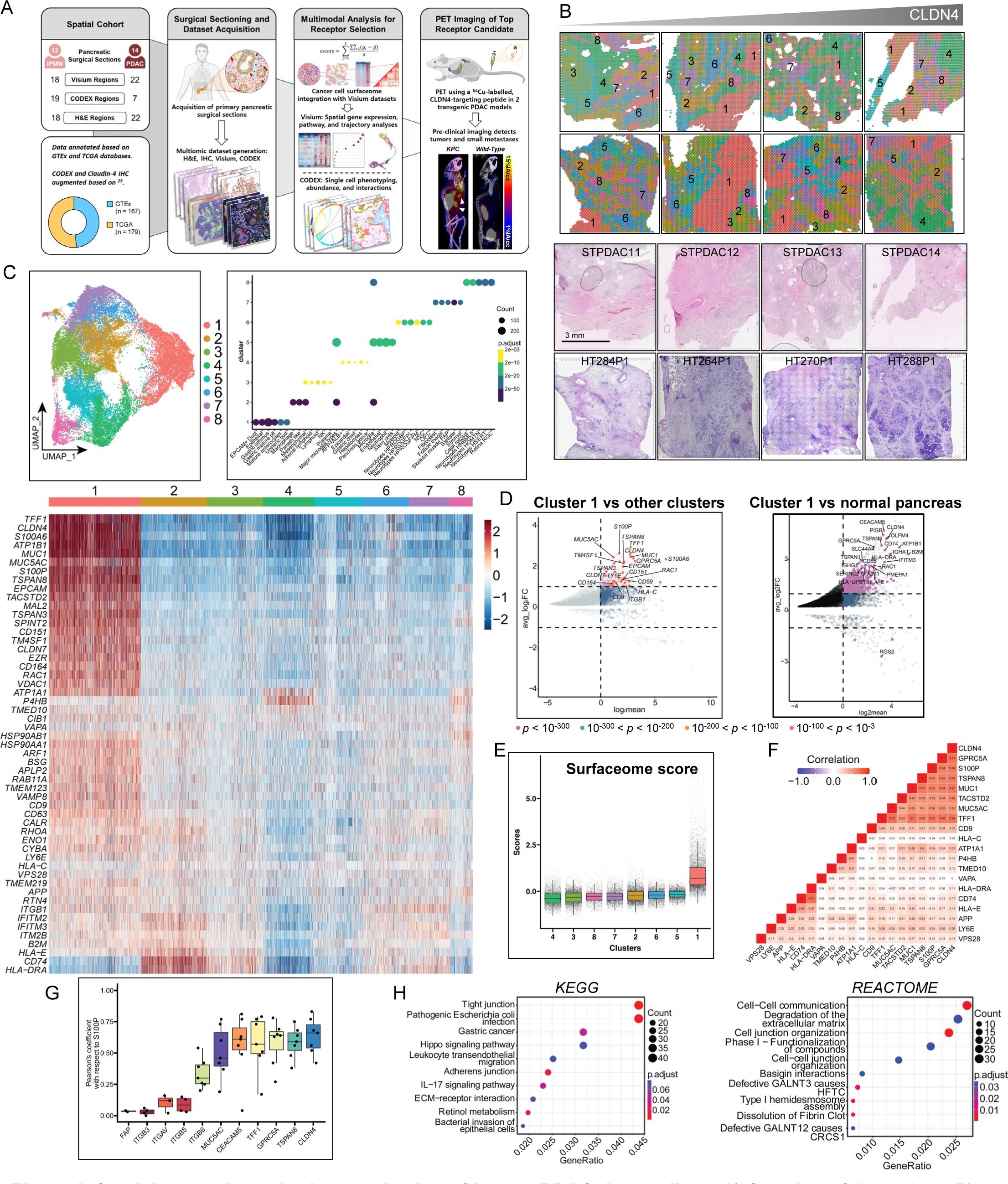
Spatial transcriptomic characterization of human PDAC tissue slices. A) Overview of the project. B) H&E of PDAC tumor excised from five patients (bottom two rows) and their corresponding spatial transcriptomic spots (top two rows) arranged based on increasing *CLDN4* expression. The spatial transcriptomes between the PDAC slices were integrated based on canonical correlation analysis (Seurat), clustered and projected on the Uniform Manifold Approximation and Projection (UMAP) dimension in B) and projected on to their histology slices. Scale bar = 3 mm. C) Leiden clustering of PDAC transcriptome projected on UMAP space and heatmap of Leiden clusters with genes selected from the PDAC data set of the cancer cell surfaceome with a specification of expression on at least 80% of cells. Cluster identities were based on gene enrichment analysis with respect to the molecular signature database. D) Differential expression of Cluster 1 versus all remaining clusters and versus samples of unaffected pancreas surrounding precancerous IPMN. E) Cancer surfaceome score based on average normalized and scaled gene expression of all cells and all cancer cell surfaceome pancreatic cancer genes in each cluster. F) Pearson’s correlation similarity matrix of select cancer cell surfaceome genes based on the PDAC spatial transcriptome. G) Pearson’s correlation of all spots for *S100P* vs key markers of interest. H) KEGG and REACTOME pathway enrichment of Cluster 1.

S100P, a largely cytoplasmic protein, was used for comparison of the spatial distribution and expression of surfaceome markers as S100P expression is absent in normal pancreas, and a specific increase in S100P expression occurs only in the PDAC tumor epithelium ^25^. S100P expression level in PDAC and IPMN is significantly higher than in nontumor pancreatic tissues, and its expression level increases with the grade of PanIN ^26^. The Pearson’s coefficient between cancer surfaceome genes and *S100P* was consistent within all spatial regions, with *CLDN4* demonstrating the highest average coefficient (Fig. 1G). The Pearson’s coefficient between *S100P* and various integrin markers or fibroblast activation protein (*FAP*) ranged between -0.3 and 0.4 (highest scoring integrins are shown in Fig. 1G) and was 0.37 and 0.01 for *ITGB6* and *FAP*, respectively. The Pearson’s coefficient between these epithelial markers and typical immune receptors (*CD3, CD8, CD64 (FCGR1A), CD14*) was also nearly zero (Supplementary Fig. S2A-B). One mechanism underlying the high correlation between enhanced *CLDN4* and selected tumor epithelial marker expression was the transcriptionally-enhanced expression of *CLDN4* with elevated RAS/PI3K/Akt expression (Supplementary Fig. S2C).

We then performed enrichment analysis (Fig. 1H) for these genes with respect to KEGG and REACTOME databases and found that Cluster 1 was enriched in tight junction (KEGG), IL-17 signaling (KEGG), and degradation of extracellular matrix (REACTOME) pathways, which implies significant stromal restructuring associated with Cluster 1.

High-resolution IHC indicated that CLDN4 protein expression was enhanced on hyperproliferative cells with a morphology consistent with cancer (Fig. 2A) (small to medium-sized irregular glands with nuclear atypia lacking tight junctions and embedded in a desmoplastic stroma). Proliferation (increased Ki67 expression) was enhanced in high-grade PanINs and cancerous ducts (Fig. 2B). Alignment of hematoxylin and eosin (H&E) images with the mRNA expression of four key surface markers (Fig. 2C) confirms enhanced *CLDN4* expression in high-grade PanINs. While mRNA expression across *S100P*, *CLDN4*, *TFF1* and *CEACAM5* was highly correlated, expression on low-grade PanINs varied with phenotype (Fig. 2C). As shown in Fig. 2A and D, in addition to the upregulation of *CLDN4* in the cancer-associated cluster, molecular imaging of CLDN4 is attractive as the tight junction protein is more accessible on diseased cells (Fig. 2D). Taken together, the results suggest that *CLDN4* is consistently elevated in PDAC and is highly spatially correlated with cancer-associated markers. Alternatively, fibroblast, integrin and immune receptor markers were not spatially correlated with *S100P* or cancer atlas markers.

**Figure 2.**
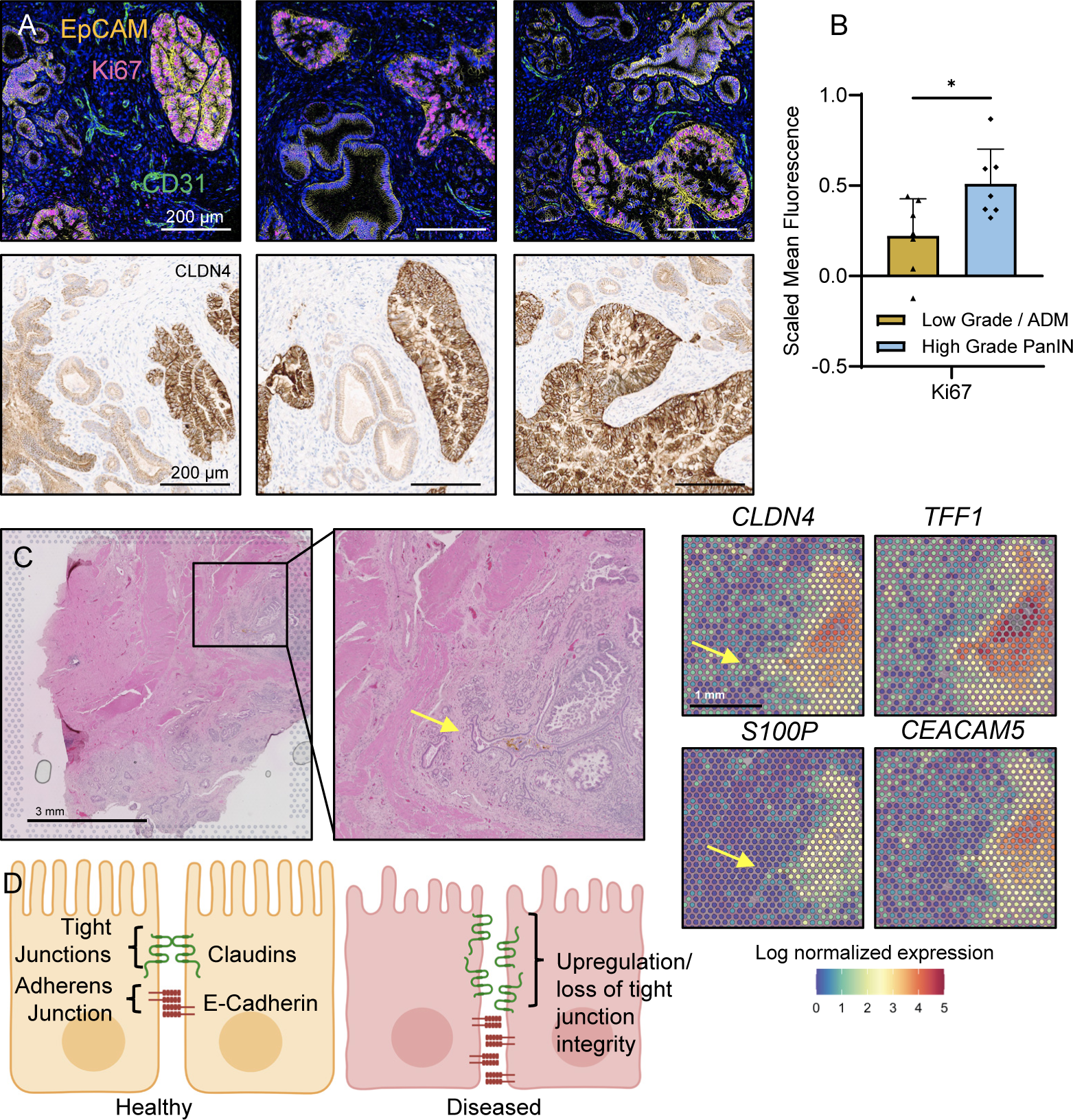
High-resolution protein imaging confirms selective expression of CLDN4 on high-grade PanINs and cancerous ducts. A) Spatial proteomic imaging of EpCAM, Ki67 and CD31 as compared with CLDN4 immunohistochemistry (IHC) on nearby sections. B) Ki67 quantification on high-grade PanIN vs low-grade PanIN/acinar-to-ductal metaplasia (ADM), C) High-resolution hematoxylin and eosin (H&E) imaging compared with mRNA expression of *CLDN4*, *TFF1*, *S100P* and *CEACAM5* in the same region validates similar expression. Yellow arrows indicate small differences in low-grade lesions. D) Illustrated summary of the changes in CLDN4 expression with disease. Significance analyzed with unpaired t-test, * = p <0.05.

### Key tumor epithelial marker expression is spatially correlated

We next visualized the spatial correlation between the top epithelial markers in PDAC and IPMNs and compared these maps with those relevant for islets and fibroblasts, where rows represent individual fields from a single patient sample (Fig. 3A). On the 100-µm spatial scale of spatial transcriptomic slides, we found that *S100P, MUC5AC, TFF1* and *CLDN4* were spatially correlated and moreover, in regions where ducts were filled with large numbers of highly-proliferative cancer cells (yellow arrow), *TFF1* and *CLDN4* were highly expressed. *ITGB6* and fibroblast activation protein (*FAP*) expression was elevated to a lesser extent in the tumor, yet on the scale of spatial transcriptomic imaging, was not enhanced near the ducts (Fig. 3A). Mapping of insulin (*INS*) expression provides an alternative view of the functional islets with *INS* expression and was typically enhanced in spots that do not express the cancer surface markers.

**Figure 3.**
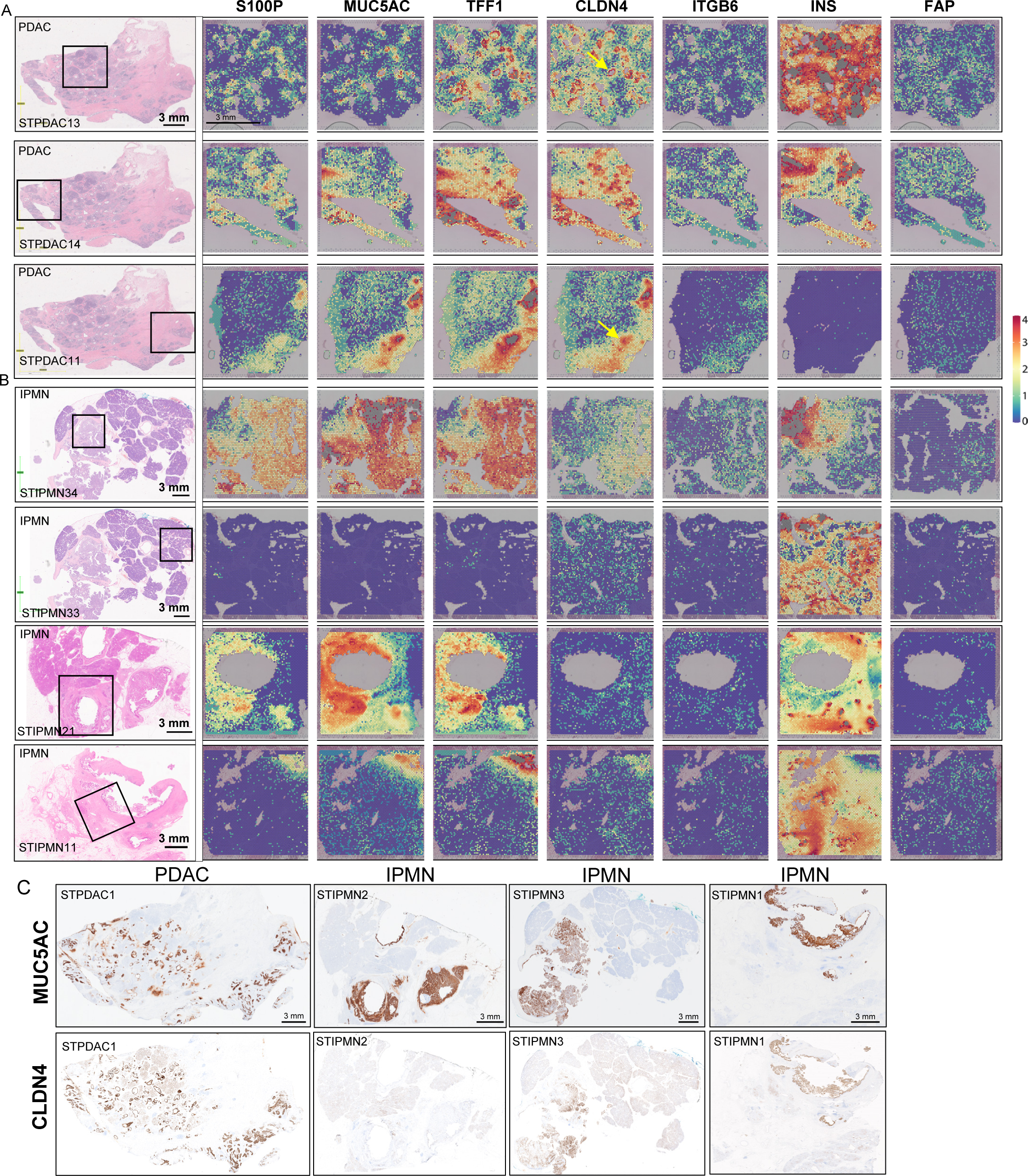
Spatial gene expression maps in PDAC and IPMN. A) Spatial gene expression maps of *S100P*, *MUC5AC, TFF1, CLDN4, ITGB6, INS and FAP* in PDAC from multiple fields in a representative patient. The H&E slides in the first column illustrate the specific location of the spatial gene expression slice on the excised tissue. Yellow arrows indicate ducts. B) Spatial gene expression levels of *S100P, MUC5AC, TFF1, CLDN4, ITGB6 INS,* and *FAP* in IPMN (tissue samples from 3 patients). C) IHC of MUC5AC and CLDN4 protein for four tissue samples (one PDAC and three IPMN).

### In IPMN samples, *CLDN4* expression was enhanced in high-grade lesions

By comparison, within the IPMNs (Fig. 3B), expression of *MUC5AC* and *TFF1* was greatly enhanced with a variable level of expression of *CLDN4* and *ITGB6*. *MUC5AC* expression was extensive in IPMNs, whereas *CLDN4* was preferentially enhanced in PDAC (Fig. 3A). Immunohistochemistry (IHC) for MUC5AC and CLDN4 confirmed that their protein expression patterns were closely correlated with their mRNA expression (Fig. 3C).

In IPMN lesions (Fig. 4A-D), cancer surfaceome genes were elevated in Clusters 6, 9, 16, and 12, with a greater fraction enhanced in Cluster 12 (Fig. 4E-F). NKX6-2^20^ mRNA counts averaged 0.002 across all Stanford IPMN samples except for STA3IPMN (average of 1.7). When compared with underlying histology, Clusters 6 and 12 were spatially correlated with elongated ducts and hyperplastic islets. We further explored Clusters 6 and 12 by differential expression analysis. Cluster 12 included elevated expression of *CLDN4* and *GPRC5A* (Fig. 4F-G, Supplementary Fig. S2C) whereas in Cluster 6, mucin producing genes, such as *MUC5AC,* were elevated. The cancer surfaceome score increased gradually across the clusters (Fig. 4H), reaching a mean score in Cluster 12 that is similar to the score observed in PDAC clusters.

**Figure 4.**
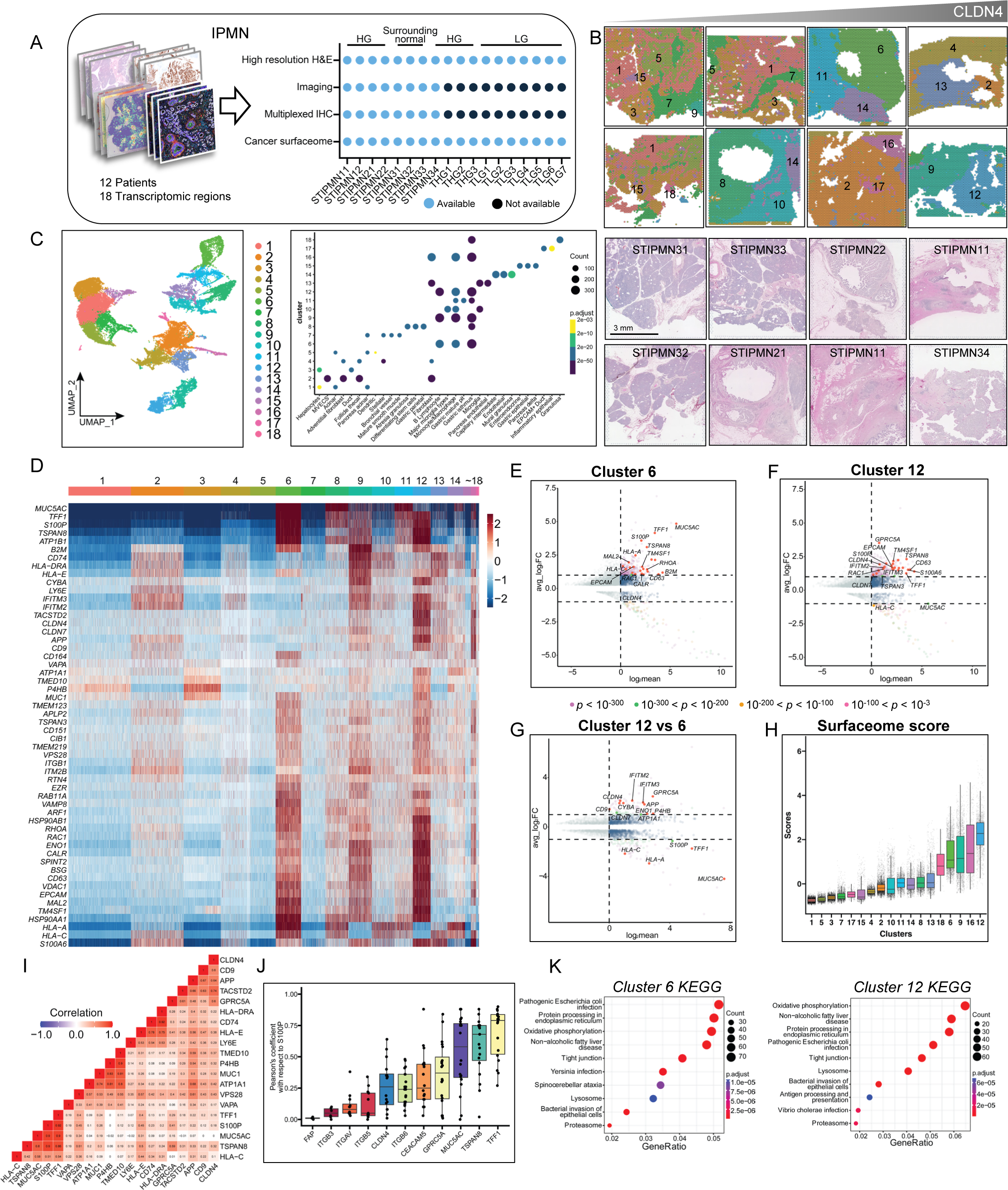
Spatial transcriptomic characterization of human IPMN tissue slices. A) Summary of the IPMN study. B) H&E slices of IPMN tumor excised from three patients (bottom two rows) and their corresponding spatial transcriptomic spots (top two rows) arranged based on increasing *CLDN4* expression. The spatial transcriptomes between the IPMN slices were merged, clustered and projected on the Uniform Manifold Approximation and Projection (UMAP) dimension in B) and projected to their H&E slices. Scale bar = 3 mm. C) Leiden clustering of IPMN transcriptome projected on UMAP space. Cluster identities were based on gene enrichment analysis with respect to molecular signature database. D) Heatmap of Leiden clusters with genes selected from the PDAC cell surfaceome atlas with greater than 80% expression across sequenced PDAC cells. E-G) Differential expression of Cluster 6 (E) and Cluster 12 (F) versus all remaining clusters and G) Cluster 12 versus Cluster 6. H) Cancer surfaceome score based on average normalized and scaled gene expression of all cells and all cancer cell surfaceome PDAC genes in each cluster. I) Pearson’s correlation similarity matrix of select cancer cell surfaceome genes based on the IPMN spatial transcriptome. J) Pearson’s correlation of all spots for *S100P* vs key markers of interest. K) KEGG pathway enrichment of Cluster 6 and Cluster 12.

In the IPMN lesions, the spatial correlation of the surfaceome markers included two groups with high inter-marker correlation, including *TSPAN8, MUC5AC, S100P and TFF1* in one group (>0.9), and *CLDN4, CD9* and *APP* in a second group (>0.6) (Fig. 4I). *TFF1* has the highest correlation with *S100P* of all markers in the IPMNs (Fig. 4J), with greater variability in the PDAC markers from Fig. 1, resulting from the range of phenotypes from normal tissue to invasive cancer observed within the cluster. Across the IPMN fields, Cluster 6 and 12 were both enriched in tight junction and infection-related pathways on KEGG enrichment analysis (Fig. 4K).

Developing an understanding of early changes in precancers could also benefit from spatial sequencing. For example, in airways, *IL-17RA* and *IL-17RC* are associated with enhanced *MUC5AC* expression ^27^. In the precancers studied here with the highest levels of *MUC5AC* expression, *MUC5AC* and *IL-17RE* expression were spatially correlated (Supplementary Fig. S2D). Evaluation of IL-17C/RE signaling in early pancreatic disease could be further probed in future work ^27,28^.

Standard medical imaging of these cases with computed tomography (CT) or magnetic resonance imaging (MRI) was also obtained and adds support for the need for molecular imaging to characterize lesions (Supplementary Fig. S3A-F). For example, retrospective analysis of contrast-enhanced MRI of the CLDN4^+^ region in lesion STIPMN34 corresponding to Cluster 12 suggested that the lesion was suspicious for cancer based on contrast enhancement (Supplementary Fig. S3G-J); however, this small region of enhancement had not previously been reported. This suggests that molecular imaging can enhance the detection of small regions with cancerous phenotypes.

### *CLDN4* increased at the pseudotime value associated with invasive disease on pseudotime analysis

Through pseudotime analysis of an epithelial-only dataset including normal pancreas and IPMN and PDAC regions, we then explored two trajectories which outline IPMN and PDAC sample types. In IPMN, the pseudotime trajectory involved an acinar-ductal transition that progressed to either IPMN or PDAC (Fig. 5A-B). By contrast, in the PDAC clusters, acinar-ductal metaplastic clusters rapidly progress to PDAC. When we further remapped the pseudotime trajectory into the spatial dimension on a unified scale, regions with a cancer-like or normal phenotype were segmented in IPMN (Fig. 5C-D). In PDAC, we found that ductal-originated cancerous cells were scattered throughout the specimen (Fig. 5E).

**Figure 5.**
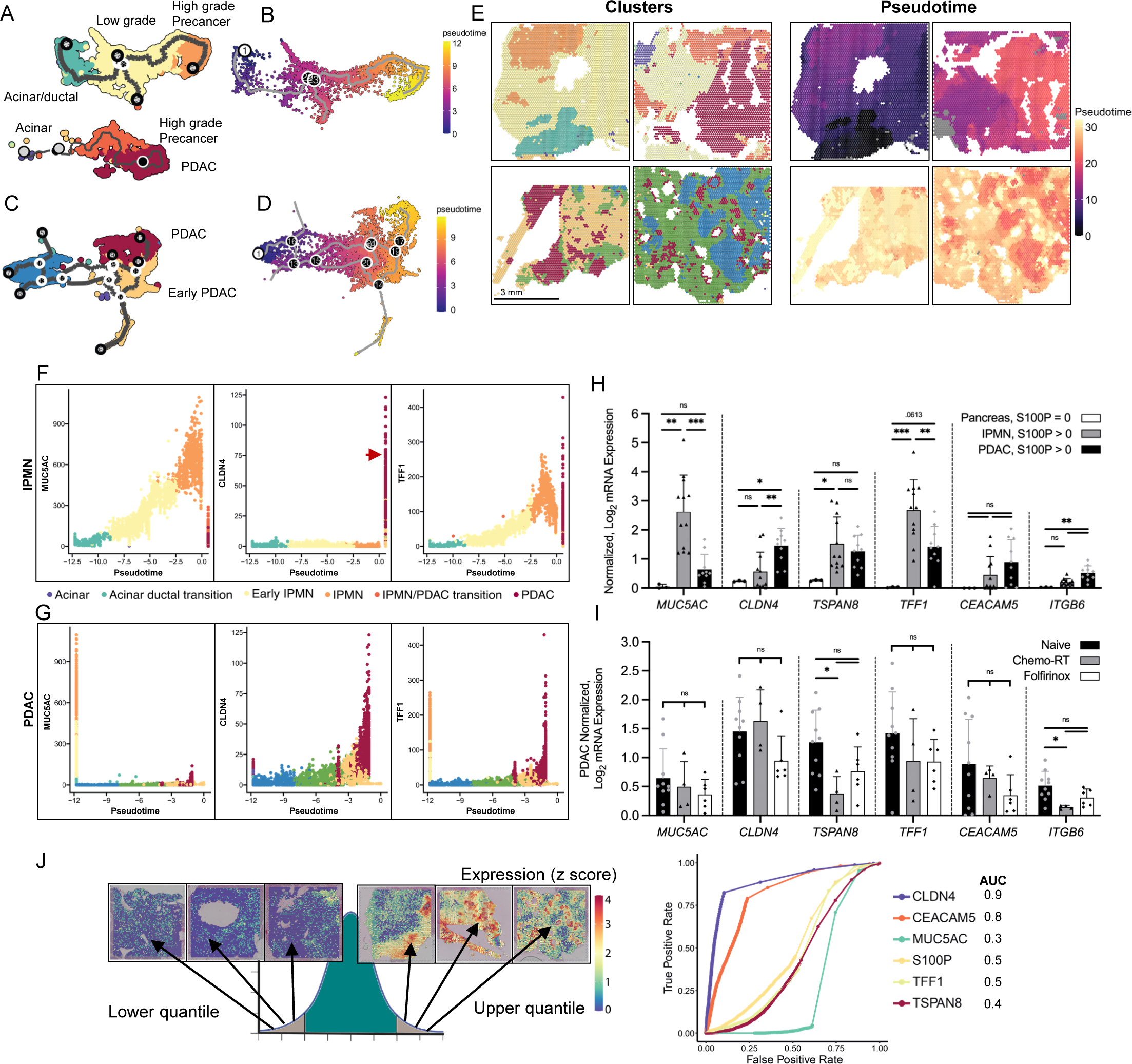
Pseudotime analysis of PDAC and IPMN human tissue spatial transcriptomes. A) Annotated spot population of clusters containing acinar ductal transition that progressed to IPMN. B) Pseudotime trajectory of acinar ductal transition to IPMN in A). C) Annotated spot population of clusters containing acinar ductal transition to PDAC. D) Pseudotime trajectory of acinar ductal transition to PDAC spots in C). E) Spot clusters projected on their corresponding histology slices and the pseudotime trajectory values projected into the spatial dimension. F-G) *MUC5AC, CLDN4, and TFF1* expression levels with respect to pseudotime in (F) IPMN and (G) PDAC, based on a pseudotime scale where 0 is invasive cancer (red arrow). H) Spots with normalized log base 2 expression of *MUC5AC, CLDN4, TSPAN8, TFF1, CEACAM5* and *ITGB6*, based on co-expression levels of *S100P* in surrounding pancreas (normal tissue), IPMN, and PDAC tissue. ns = not significant. I) Spots with identical genes and expression level based on co-expression levels of *S100P* in naïve PDAC tissue, PDAC tissue from patients treated with chemotherapy and radiation therapy (Chemo-RT), and PDAC from patients treated with chemotherapy Folforinox. J) Receiver operating curve obtained from mRNA spatial analysis of all PDAC and IPMN samples for surfaceome genes. Statistics were analyzed with one-way ANOVA for each column, * = p <0.05, ** = P <0.005, *** = P < 0.0005, **** = p < 0.00005 unless otherwise stated.

The IPMNs highly expressed *MUC5AC* and *TFF1* at earlier pseudotime values, with *MUC5AC* expression decreasing over time (Fig. 5F-G). Alternatively, *CLDN4* increased at the pseudotime value associated with invasive disease (Fig. 5F-G), confirming the results from Fig. 4G, which also demonstrated that *CLDN4* was enhanced in the IPMN region with the highest cancer surfaceome score.

The enhanced expression of *CLDN4*, *TSPAN8,* and *TFF1* on cancer cells was not significantly reduced after treatment with chemotherapy with or without radiation (Fig. 5H-I). The area under the curve (AUC) for the differentiation of PDAC from normal or precancerous tissue for *CLDN4*, *S100P*, *MUC5AC*, *TFF1*, *CEACAM5*, and *TSPAN8* was 0.9, 0.5, 0.3, 0.5, 0.8, and 0.4, respectively (Fig. 5J).

### Activated fibroblasts were organized in thin layers surrounding high grade lesions

We then performed 51-plexed cytometric imaging via codetection by indexing and immunohistochemistry (Fig. 6A). With this enhanced resolution, ductal lobules on the order of tens of microns can be visualized and the phenotype determined (Supplementary Fig. S4). In regions harboring high numbers of high-grade PanINs (large CD66^+^ ductal features) (Fig. 6B), CD45^+^ immune infiltration, ADM (small Keratin8/18^hi^EpCAM^+^ ductal features), desmoplasia (a- SMA^+^Podoplanin^+^ myofibroblasts), and overall loss of acinar structure (EpCAM^+^ epithelial cells) was observed. Importantly, while activated fibroblast markers were not detected via spatial transcriptomics, thin bands of activated fibroblasts (a-SMA^+^Podoplanin^+^ myofibroblasts) were visualized at single cell level surrounding ducts.

**Figure 6.**
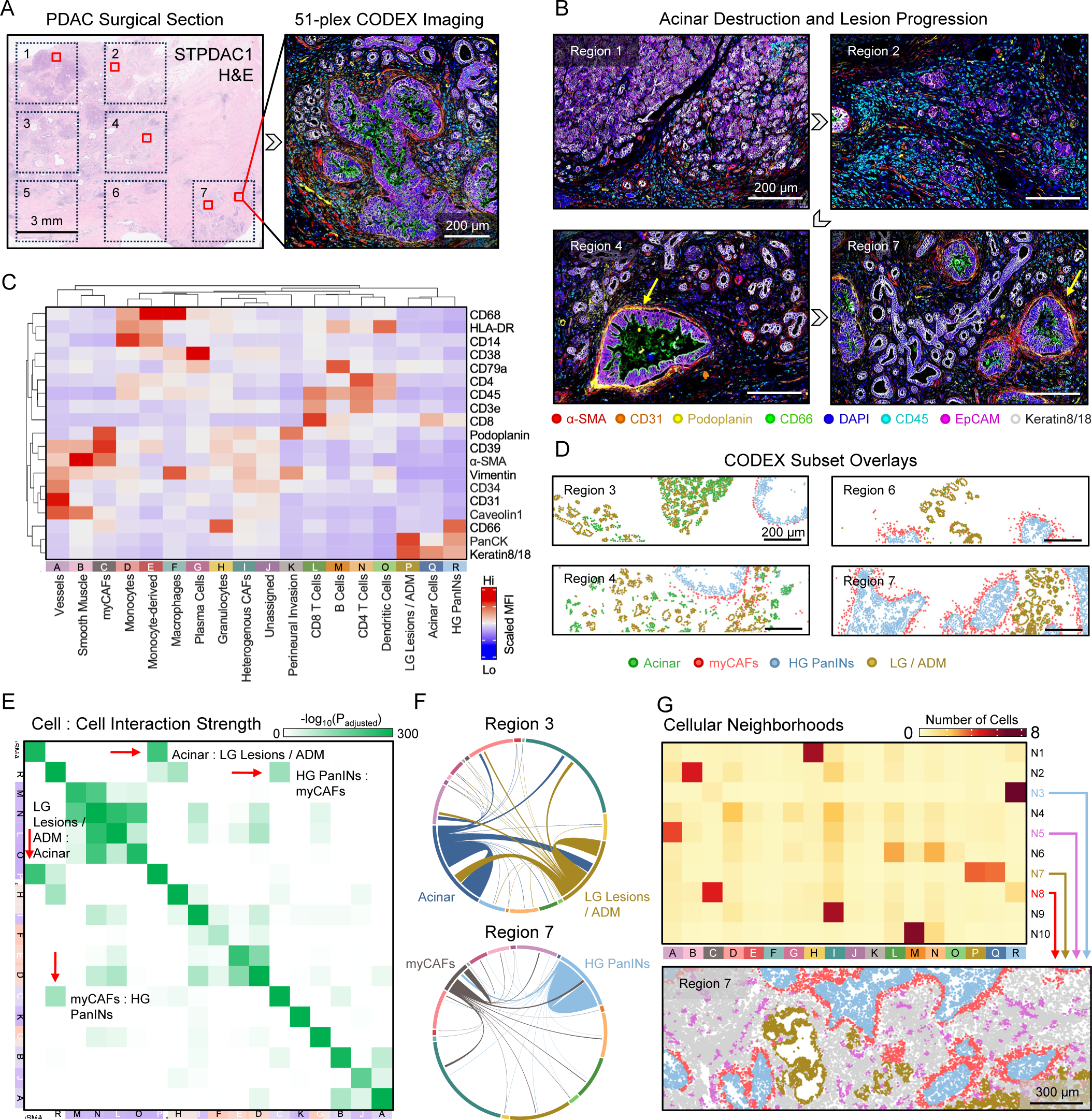
Mapping of protein at single cell level in PDAC. A) 7 regions obtained from a single PDAC surgical section were stained with a 51-plex CODEX panel. B) Representative CODEX images depicting acinar destruction, immune infiltration, and myCAF stratification in regions with increasingly severe levels of disease progression (grey arrows). Fibroblasts encompassing cancerous duct shown with yellow arrow. C) Scaled expression heatmap for 18 manually annotated cell clusters that were generated via unsupervised Leiden clustering. D) “Acinar”, “myCAF”, “High grade (HG) PanINs”, and “Low grade (LG)/ ADM” populations were plotted across several regions, highlighting discrete spatial organization of these cell types. E) Cell-cell interactions (pairwise adjacency) were quantified for all 18 clusters, and select interactions are designated with red arrows (Acinar : Low grade / ADM and myCAFs : High grade PanINs). F) Chord diagrams of regions 3 and 7 highlight heterogeneity in the frequency of cell-cell interactions (width of chord) and individual populations (length of arc). G) 10 neighborhoods (k), each composed of 10 cells (knn), were generated for the entire CODEX dataset. The abundance heatmap shows the average cellular composition of each neighborhood across all regions.

To further dissect individual cell phenotypes, we performed unsupervised clustering and manually annotated cells based on canonical expression profiles for leukocytes, endothelial cells, and epithelial cells (Figure 6C). By extracting the Acinar, myCAF, High Grade PanIN, and Low Grade PanIN/ ADM populations as annotated in Fig. 6C, we found across all CODEX regions that myCAFs stratified ducts containing highly proliferative tumor cells (Fig. 6D). These findings were validated by calculating the likelihood, or strength, of cell-cell pairwise interactions for all regions, showing that acinar cells were likely to interact with ADM, and myCAFs, with high grade PanINs (Fig. 6E). We further visualize these cell-cell contacts using chord diagrams, emphasizing the connection of myCAFs and high grade PanINs (Fig. 6F).

High-grade PanIN cells and myCAFs each comprised unique neighborhoods that were well-defined and co-localized spatially (Fig. 6G, Neighborhoods 3 and 8, respectively). Early lesion cells, ADM, and acinar cells were grouped together into neighborhood 7, supporting the cell-cell interaction results from Fig. 6H and likely representing areas of tissue where early pancreatic oncogenesis is likely present. Neighborhood 5 was built primarily of CD31^+^Caveolin1^hi^ endothelial cells and a few heterogeneous CAFs (stellate cells are likely found in this cluster), reflecting angiogenic neo vessels distributed across the tumor landscape.

As previously shown in Fig. 2B, high-grade PanINs expressed high levels of *CLDN4* across the entire duct. CODEX results indicate that myCAFs were organized in thin surrounding layers. The relative spatial extent of expression and the separation between myCAFs and cancerous epithelial cells further motivate the selection of *CLDN4* as an imaging and therapy target.

### Tertiary lymphoid organs and sparse immune infiltrates reveal leukocyte heterogeneity

As we considered possible molecular-imaging strategies, we also assessed the distribution of immune cells within PDAC and precancers. Immune cell density was heterogeneous due to the presence of both scattered individual leukocytes and tertiary lymphoid structures and organs (TLOs). Both proliferation (Ki67) and immune cell density were enhanced in the IPMN center (Supplementary Fig. S5A-D). TLOs were distributed throughout IPMNs/PDACs and detectable on spatial transcriptomics due to their millimeter-scale sizes, spherical/oblong structures, and intense expression of immune-related genes (Supplementary Fig. S6A-B). TLO differentially-expressed genes (DEGs) and associated pathways were enriched for B/T cells, B-cell receptor/T-cell receptor (BCR/TCR) signaling, chemokine/cytokine interactions, immune cell activation, proliferation, and differentiation (Supplementary Fig. S6C-F). The spatial organization of these structures was well-defined, as T cells (CD4 and CD8), B cells (CD20 and CD21), dendritic cells (CD141), plasma cells (CD38) and high endothelial vessels (CD31) were shown to occupy specialized zones within TLOs (Supplementary Fig. S6G). Upon filtering the CODEX TLO data for T and B cells, we also found that CD45RO and CD44 were highly co-expressed on both CD4 and CD8 T cells, indicative of an activated memory phenotype after antigen exposure (Supplementary Fig. S6H-I). Furthermore, Ki67 was primarily found on TLO-core B cells, indicating somatic hypermutation and protective antibody production (Supplementary Fig. S6H- I).

Within the IPMNs (Supplementary Fig. S7A-D), lymphoid structures were frequent; however, these structures often lacked the organization demonstrated in mature TLOs (Supplementary Fig. S7A-B). We then quantified the leukocyte frequency and phenotype across the IPMN and treatment-naïve PDAC regions (Supplementary Fig. S7C-D). Leukocyte frequency increased in both IPMN and PDAC tissues compared with surrounding pancreas (Supplementary Fig. S7C).

The high immune cell variability in the IPMN results from a wide range of immune infiltration (20 to 60% of total cells) within each acquired region (Supplementary Fig. S7C).

### Results point to biologically-driven molecular imaging of the tight junction protein CLDN4

Based on our spatial multi-omic analyses, CLDN4 emerged as an imaging and potential therapeutic target due to its high differential expression between high-grade lesions and normal tissues, consistent correlation across mRNA expression with other cancer markers such as *S100P* and late-stage expression in pseudotime. Small peptides are superior for targeting due to the ease of synthesis, comparable potential affinity and specificity, improved pharmacokinetic profiles, and low immunogenicity. Clostridium perfringens enterotoxin (Cpe), a natural ligand to CLDN4, is 35 kDa. The C-terminus Cpe (cCpe), a truncated form of Cpe, binds to the extracellular segments of CLDN4 with a nanomolar binding affinity (KD) ^29^. Several short peptides that bind to the ECL4 loop in CLDN4 have been identified ^30^. Our ligand validation with docking simulation in human CLDN4 (Supplementary Fig. S8A) suggests that Cpe30 MT2 (NSSYSGNYYSIL, KD of

1.97 nM, sequence will be abbreviated to C4BP (claudin-4 binding peptide) hereafter) is promising for human imaging. We therefore synthesized DOTA-PEG1-C4BP, mimicking the native sequence of NSSYSGNYYSIL, on a rink amide resin using a microwave assisted peptide synthesizer. DOTA, a Cu-64 chelator, which also incorporates various therapeutic and diagnostic metal isotopes, was attached after one PEG spacer unit to avoid steric hindrance (Supplementary Fig. S8B). High-performance liquid chromatography (HPLC) purification afforded DOTA-PEG1-C4BP in >95% purity with exact mass (M+H^+^, calculated 1897.9 Da, found 1898.0 Da) and confirmed with matrix-assisted laser desorption/ionization (MALDI) mass spectrometry (Supplementary Fig. S8C). The radiolabeling of DOTA-PEG1-C4BP (2 nmol) with ^64^CuCl^2^ (70 MBq) conferred ^64^Cu- DOTA-PEG1-C4BP (28 MBq/nmol, ^64^Cu labelled DOTA-PEG1-C4BP will be abbreviated to ^64^Cu- C4BP here after) in 80% non-decay corrected yield with >99% radiochemical purity (Supplementary Fig. S8D).

Imaging of *Kras^LSL-G12D/+^;Rosa26^LSL-tdTomato/LSL-tdTomato^(KT);Ptf1a^CreER^;Trp53^fl/fl^*mice was then performed at 49, 84, and 120 days after tamoxifen injection (Fig. 7A). In this model, substantial CLDN4 protein expression was detected in pancreatic tumors and metastases (Fig. 7B). At 49 days post tamoxifen treatment, a dynamic PET/CT scan was simultaneously acquired with tail vein administration of ^64^Cu-C4BP (n=4, ∼0.1 μg/mouse), and imaging in the 25-30 minute window displayed a significant retention of radiolabeled ligand at the tumor sites in this transgenic mouse model (white arrowheads, Fig. 7C). Maximum intensity projection (MIP) and slice imaging detected the presence of tumors and metastases only in cancer-burdened mice and not in wild-type animals (Fig. 7C-D). Image-based quantification assessed the accumulation of injected activity per cc (%IA/cc) in the tumor and blood, peaking at ∼15%IA/cc (2 min) and clearing to < 5%IA/cc (30 min), respectively, in tumor-burdened mice 49 days after tamoxifen administration (Fig. 7E). At later tamoxifen time points, accumulation had increased to ∼25%IA/cc in metastases and was retained at ∼18%IA/cc in the pancreas within 30 minutes after ^64^Cu-C4BP injection (Fig. 7F). Since MRI provides more detailed soft tissue anatomical reference than CT and KPC mouse models have been shown to have increased signal intensity at the tumor location in T2-weighted MRI, we performed multi-modal imaging including T2-weighted MRI, PET/CT, and bright field imaging to validate that the region retaining ^64^Cu-C4BP signal was consistent with pancreatic cancer (Fig. 7G). *Ex vivo* PET imaging confirmed the localized accumulation of ^64^Cu-C4BP within the colon, pancreas and liver in transgenic mice (Fig. 7H). Biodistribution showed 6.6 ± 3.9 %IA/g in the kidneys and 7.7 ± 5.4%IA/g in the liver at 22 hours post injection (Supplementary Table S3). Additionally, the pancreas-to-muscle ratio in transgenic mice (3.71 ± 0.56%IA/g, *p*=0.003) was significantly higher than age-matched wild-type mice (1.53 ± 0.1%IA/g) (Supplementary Table S3).

**Figure 7.**
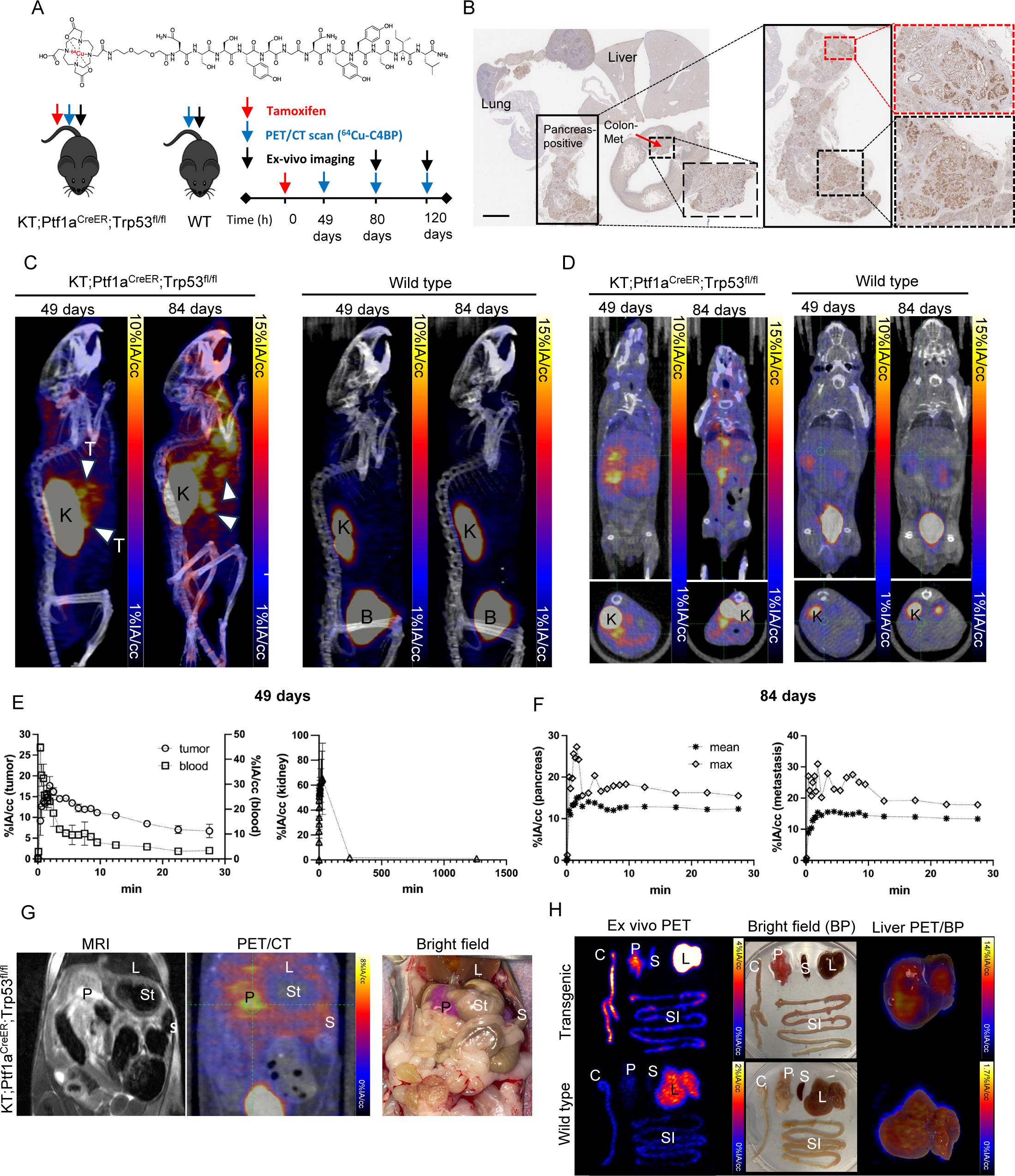
Molecular imaging of claudin-4 (CLDN4) in a PDAC mouse model. A) CLDN4-binding peptide and imaging scheme for study in tamoxifen-induced pancreatic cancer in the *-KT;Ptf1a*^CreER^*; Trp53*^fl/fl^ mouse model of PDAC. B) CLDN4 immunohistochemistry of organs (liver, lung, colon and pancreas) from the tamoxifen-induced PDAC mouse model. We used ^64^Cu-C4BP and demonstrated tumor-specific accumulation. This transgenic model expresses CLDN4 in both the pancreatic lesions and metastases (Met) after tumor induction. Scale bar: 2 mm. C) PET/CT maximum intensity projection (MIP) images acquired in the first 30 minutes after ^64^Cu-C4BP injection in the transgenic mouse model at 49 or 84 days after tamoxifen treatment. White arrowheads indicate accumulation of ^64^Cu-C4BP at the tumor sites. D) PET/CT slice images showing retention of ^64^Cu-C4BP within the pancreas (crosshair). E) Time-activity curves (TACs) for blood, tumor, and kidney at 49 days (note that ^64^Cu-C4BP clears from the kidney). F) TACs for the pancreas and metastases at 84 days. G) Magnetic resonance imaging (MRI), PET/CT, and bright field images of abdominal region from the same mouse at 120 days after tamoxifen treatment. ^64^Cu- CBP in the PET/CT image is well localized to the pancreas in the MRI and bright field images. H) In the same model at ∼84 days after tamoxifen treatment, *ex vivo* PET imaging with ^64^Cu-C4BP detects disease in the pancreas, colon and liver. Tumor cells express tdTomato in this model producing a red hue in the pancreas and colon. Liver accumulation is high and associated with metastasis in the transgenic model (note scale). A smaller amount of activity accumulates in the liver due to clearance in the wild-type (WT) mouse. Abbreviations: L: liver, P: pancreas, St: stomach, S: spleen, C: colon, SI: small intestine, K: kidney, B: bladder, T: tumor.

We then repeated this imaging study in small (5 mm) tumors in the Capan-1 xenograft mouse model (Supplementary Fig. S9), which overexpresses human CLDN4 (Supplementary Fig. S9A- B) to a greater extent than alternative pancreatic models. Human CLDN4 is highly conserved as compared to mouse protein. Accumulation was again verified by PET MIP and slice imaging at multiple time points (Supplementary Fig. S9C-D), with a retention of ^64^Cu-DOTA-C4BP in the tumor sites (indicated by white arrowheads) as a function of time. A blocking study achieved by co-injection of a 1000-fold excess of the cold compound impeded 54% (*p*=0.0003) of ^64^Cu-DOTA- C4BP accumulation at the tumor (Supplementary Fig. S9C-D, G). A fraction of the activity continued to circulate in the blood for more than 30 min (Supplementary Fig. S9E-F). Tumor accumulation reached ∼6.9%IA/cc at early time points (Supplementary Fig. S9G), and radioactivity rapidly cleared from the kidneys (Supplementary Fig. S9H).

We next explored the spatial extent of a therapeutic radiation dose delivered to the human surgical sample lesions, assuming that a therapeutic radiopharmaceutical successfully bound in a manner proportional to the local mRNA concentration (Supplementary Fig. S10). We compared the distribution of *CLDN4* and *MUC5AC* mRNA to that of *S100P* as the gold standard (Supplementary Fig. S10A), where red spots would represent regions with *S100P* above baseline but in the absence of the CLDN4 or MUC5AC targeted drug. We found that nearly all of the 100 µm spots which express *S100P* also express *CLDN4* (Supplementary Fig. S10B). Furthermore, given the effective therapeutic range of alpha or beta emitters (Supplementary Fig. S10C-D), few voxels, if any, would be excluded from a therapeutic dose using a CLDN4-targeted radiopharmaceutical. This finding will be assessed *in vivo* in future work.

Finally, we analyzed subclusters of the cancer-associated Cluster 1 in Fig. 1 (Supplementary Fig. S11) in order to characterize the phenotype of spots without *CLDN4* overexpression. *CLDN4* expression was not enhanced in Subcluster 10 (Supplementary Fig. S11A-B), and differential expression analysis demonstrated enhanced expression of *TFF2, MUC6, LYZ, NKX6-2, PGC, GKN2, VSIG1, CXCL17, CLDN18 and GOLM1* in this subcluster (Supplementary Table S4 and Fig. S11C). A fraction of these spots had a high Pearson spatial correlation between *TFF2, MUC6 and LYZ* overexpression, corresponding to mucous neck cells or antral gland cells ^31^ (Supplementary Table S4 and Fig. S11D). In addition, in a fraction of spots, *NKX6-2, PGC, GKN2, VSIG1, CXCL17 and GOLM1* expression was spatially correlated with a Pearson correlation greater than 0.4 (Fig. S11D). Overexpressed genes in each of these correlated regions were enhanced earlier in pseudotime as compared with *CLDN4* (Fig. 5F, Supplementary Fig. S11E), suggesting that these spots correspond to a lower grade phenotype.

## Discussion

The development of biologically-inspired strategies to image and treat cancers with precise molecular targeting has high significance. With the emergence of spatial methods and data sets, there is an opportunity to use molecular imaging both to detect cancer and also to design and implement therapeutic strategies. Here, for the first time, we applied spatial -omics techniques to characterize the distribution of cancer cell surface markers in order to design an imaging and theranostic strategy. We created a cancer surfacesome score and found that high scores were spatially correlated with a cellular morphology consistent with cancer.

Most importantly, we found that the expression of a small number of tumor surface markers, including *CLDN4*, was consistent across the patients studied, with less variability than integrin, fibroblast or immune cell markers. In the cancer cluster, the expression of surface markers *CLDN4*, *GPRC5A*, *TSPAN8*, and *CEACAM5* was enhanced more than 13-fold compared with the normal pancreas, whereas *FAP* and *ITGB6* were differentially expressed by less than 4-fold. *CLDN4* was expressed on ∼88.8% of PDAC cells in ^11^, and here we found that ∼89% of spots in the cancer-associated cluster expressed elevated *CLDN4*. The fraction of spots overexpressing CLDN4 increased to 97.5% upon integration (as compared with merging) of the data, reflecting differences in the sequencing depth at different sites. Such differences should be further analyzed in the future. Receiver operating characteristic analysis of PDAC diagnosis based on the top surface markers indicated that the area under the curve (AUC) was greatest for *CLDN4* (∼0.9).

Enhanced *CLDN4* expression and localization in regions of invasive cancer were consistent throughout PDAC lesions whether treatment naïve or post therapy. *CLDN4* expression increased later in the transition to invasive cancer than *MUC5AC* or *TFF1*, suggesting that this marker may differentiate between premalignant and invasive lesions. A retrospective evaluation of contrast MRI imaging suggested that molecular imaging of CLDN4 could have been useful in detecting a region with a high cancer surfaceome score. Based on the analysis of IPMN lesions in ^20^, in spots with minimal *NKX6-2* expression, *CLDN4, CXCL2*, *CD74, LY6E,* and *SPP1* expression was greater and was associated with PDAC progression and neoplastic progression of IPMN ^20,32^. *NKX6-2* expression was low in our cancer cluster and the results support our pseudotime analysis and solidify *CLDN4* as a marker to differentiate high- and low-grade lesions.

The resulting PET imaging approach demonstrated exquisite sensitivity for the detection and mapping of small tumors and metastases in mouse models with molecularly-specific accumulation. While *CLND4* is expressed in the normal colon, here, we found that the resulting radiopharmaceutical accumulation in normal tissue was low, likely due to the tight junction expression of this target. Accumulation of the CLDN4-targeted peptide reached ∼25%ID/cc in metastases and ∼18% IA/cc in the pancreas at 30 minutes after ^64^Cu-C4BP injection. Preclinical single photon emission computed tomography (SPECT) imaging of CLDN4 has been explored with an antibody ^33^ and other peptides in the past ^34^; however, our ligand enhanced accumulation and improved signal-to-noise ratio. For imaging of integrins, αvβ6 has been the most promising with affinity of <55 nM and uptake of ∼2-4%ID/cc in a pancreatic cancer xenograft ^35^ or affinity of 0.047 nM and ∼8% ID/cc accumulation in lung or breast cancer models ^36^ reported. However, our results suggest that the expression and localization of integrins on PDAC cells are lower than that of *CLDN4* and are reduced by chemotherapy or chemo-radiation therapy.

A single cell-based ‘omic analysis is required to fully characterize the interaction of immune and fibroblast components with epithelial targets. The immune cell distribution within pancreatic lesions was complex and will likely prove challenging to harness as a cancer-specific diagnostic feature. Without adding CODEX, we would have missed the presence of thin layers of activated fibroblasts surrounding cancerous ducts due to the relative spatial resolution and sensitivity of current spatial transcriptomic methods. Still, the numbers of myCAFs near the cancerous ducts was small and targeting these cells may not be efficient for a theranostic strategy.

Peptide targeted radionuclide therapy (PTRT) is a promising strategy for PDAC and the CLDN4- targeted peptide described here shows promise to be incorporated in PTRT. We found that a beta radiopharmaceutical field generated from a *CLDN4*-targeted therapy would generate an approximately uniform field for the entire cancerous specimen. Systemic administration of ^177^Lu- targeted radiopharmaceuticals can result in specific targeting of cancer cells throughout the body and induces the cytotoxic effects of ionizing radiation ^37^. PTRT has improved the overall survival of neuroendocrine tumors ^38^ and metastatic castration-resistant prostate cancer ^39^, and is approved by the FDA. A future goal of this work is to incorporate CLDN4 targeting in the treatment of PDAC. Thus, the results suggested that a data-driven target selection would result in the selection of a target such as CLDN4 and we are encouraged to find that promising affinity and accumulation are feasible.

In summary, we find that spatial transcriptomics is highly useful for the selection of molecular imaging agents. While the imaging of integrins or fibroblasts can be useful, the development of epithelial-based molecular imaging and theranostic agents is desirable. CLDN4 is one such target, and imaging of this target with high sensitivity and specificity is feasible.

## Online Methods

### Spatial Transcriptomic Processing and Analysis

Formalin-fixed, paraffin-embedded (FFPE) human pancreatic cancer tissue sections were placed on spatial transcriptomic slides for spatial transcriptomic sequencing based on the 10x FFPE workflow (10x Genomics, Pleasanton, CA). Individually indexed libraries were sequenced on the Novaseq 6000 (Illumina Inc., San Diego, CA). Raw sequencing counts was processed through the SpaceRanger pipeline (10x Genomics) and aligned with a human reference (GRCh38). Downstream transcriptomic analysis was performed based on the Seurat framework with modifications. Spatial transcriptomic samples were merged or integrated using R function for merging and Seurat integration function based on canonical correlation analysis. Normalization was performed with SCTransform, which minimizes technical noise such as difference in sequencing depth. Stanford-acquired data detected an average of 3706 spots per sample and 17943 aligned gene features, while PDAC datasets from HTAN^21^ detected 4992 spots per sample with 36601 aligned gene features. IPMN datasets from NCBI GEO (GSE233254) detected an average of 941 spots with 36601 aligned gene features^20^. Principal component analysis was performed, and the elbow plot indicated that the first 75 principal components were sufficient to describe over 99% of the data.

To optimize for subsequent Leiden clustering parameters such as number of principal components, resolution, and number of k nearest neighbors, the average silhouette score of all clusters was used as a tuning metric to design our hyperparameter tuning grid. Based on our optimized parameters, we performed Leiden clustering and projected the clustering results on the Uniform Manifold Approximation and Projection (UMAP) dimensions. We then computed the top 10 differentially-expressed genes of each cluster compared to all the remaining clusters using Seurat’s built-in functions. Since in spatial transcriptomics, each spot is not a single cell, and annotation based on cell types may not be appropriate, we performed statistical overlap calculations comparing clusters to databases. Using the molecular signature database cell signature gene sets ^24^, we performed enrichment analysis using differentially expressed genes from each cluster. To plot cancer cell surfaceome gene expression levels throughout the clustered spots, we selected for surfaceome genes specific to PDAC and that are at least 80% upregulated compared to normal tissue. We further developed a cancer cell surfaceome score based on the total sum average normalized and scaled gene expression of cancer cell surfaceome in each cluster:

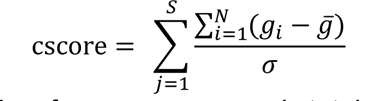

Where *S* is number of cancer cell surfaceome genes, *N* is total number of spots, *g*_*i*_ is normalized and scaled count of each gene, *g̅* is average nomalized and scaled count and σ is variance. Pearson’s correlation coefficients for each spot and each sample were calculated based on the standard Pearson’s correlation equation. Gene set analysis was performed with clusterProfiler enricher function in KEGG, GO, and REACTOME databases with top 10 most significant pathways plotted ^40^. Differential expression fold change values mapped onto KEGG pathway plots were further calculated with pathview function in R and then simplified to key pathways. Pseudotime trajectory analysis was performed with monocle3 on a merged dataset containing two surrounding normal tissue slices, two PDAC slices, and two IPMN slices. From our population annotations, we have identified two origins which we defined as acinar ductal transitions. Acinar-ductal cell transitions and metaplasia have been demonstrated by other groups as an important component in the development of PDAC ^2,41^. To focus on the development from acinar-ductal transition to full cancer, we removed cancer associated fibroblasts, plasma cells, and immune cells in our trajectory. The calculated trajectories were analyzed based on their corresponding partitions in, respectively, IPMN and PDAC. To explore the spatiotemporal relationship between PDAC phenotypes, multiple pseudotime partitions were connected sequentially, and then pseudotime values were plotted with respect to their x-y spatial coordinates.

### CODEX Processing and Analysis

Formalin-fixed paraffin embedded (FFPE) pancreas slices were stained with a 51-plexed antibody panel as in ^42^. After imaging processing, DAPI-based cell segmentation was performed with the DeepCell algorithm through the EnableMedicine portal. We further filtered the segmentation results based on size of the cell, total biomarker intensity, DNA channel intensity and signal coefficient of variation using EnableMedicine’s SpatialMap package on R. The filtered segmentation results were then normalized and scaled for principal component analysis. Unsupervised Leiden clustering was then executed to identify cell populations of interest. Populations were manually annotated using expression intensity patterns and spatial organization. Cell-cell interactions were calculated using a k nearest neighbors algorithm to create a graph network of individual cells (10 nearest neighbors were used), and then pairwise adjacency was calculated using a hypergeometric method. Neighborhoods were constructed using the previous calculated nearest neighbors graph (knn = 10), and k = 10 neighborhoods were built based on an optimal silhouette score. Segmented regions were also exported to the cloud-based cytometry platform OMIQ for additional visualization.

### Therapeutic Simulations

To model peptide binding to a given spatial transcriptomic region, we first labelled spatial transcriptomic spots using an expression threshold for *MUC5AC/CLDN4* and *S100P*. The thresholds were assigned by first selecting spots that had expression greater than 0 for *MUC5AC/CLDN4* or *S100P*, and then calculating the first quartile of expression for those genes in the corresponding positive expressing spots. Then, all spots for a spatial transcriptomic region were assigned into four groups (Gene < Threshold and *S100P* < Threshold, Gene < Threshold and *S100P* > Threshold, Gene > Threshold and *S100P* > Threshold, Gene > Threshold and *S100P* < Threshold) using the previously calculated thresholds. To model killzones for alpha and beta emitters targeted to *MUC5AC* or *CLDN4*, respectively, we first assumed a targeted peptide would have 100% binding efficiency to a target. With this assumption, expression intensity served as a proxy for particle emission density, and spatial transcriptomic spots were sized to have a radius of 60 or 600 um for an alpha or beta emitter, respectively.

### **Transgenic mouse model** ^41^

To deconstruct the requirements for pancreatic cancer development from adult pancreatic cells, the Attardi laboratory has developed a *Kras^LSL-G12D/+^;Rosa26^LSL-tdTomato/LSL-tdTomato^;Ptf1a^CreER^;Trp53^fl/fl^*mouse model^41^ by activating oncogenic Kras^G12D^ and deleting *Trp53* in mouse pancreatic cells with a tamoxifen-inducible knock-in *Ptf1a^CreER^* allele. Importantly, to mimic human PDAC development, which occurs in adults, mice are aged to adulthood (8 – 10 weeks) before tamoxifen introduction.

### Radiolabeling

All radiolabeling experiments were conducted under the Controlled Radiation Authorization (CRA) approved by Stanford University. For the radiolabeling of DOTA-PEG-C4BP with ^64^Cu (*t1/2*=12.7 h), ^64^CuCl2 (74 MBq, 2 mCi; 0.1 M HCl) from Washington University was mixed with freshly prepared labeling buffer (200 μL, 0.1 M ammonium citrate/0.1M ascorbic acid, pH 6.0). The peptide (2 nmol) was added to reach molar activity (74 MBq/nmol, 2 mCi/nmol). The reaction cocktail was then mixed at 600 rpm, 60 °C for 15 minutes, and the reaction progress checked by radio-TLC. The solid phase extraction cartridge (C18 SepPak plus light column) was pre-activated with 5 mL 99.9% ethanol followed by 10 mL water. The radiolabeled peptides were eluted from the cartridge with ∼0.5-1 mL solution prepared with 1:1 volume ratio of 99.9 % EtOH:water. The eluted solution was diluted with 0.9% sterile NaCl with 10 mg/mL ascorbic acid (pH 7) until a final ethanol concentration reached less than 10%. Radiolabeling yield and radiochemical purity were analyzed with a TLC scanner (Bioscan) and HPLC (C18 column 4.6x250 mm, Phenomenex) connected to radiodetector, respectively. Non-decay corrected isolated yield, radioTLC yield, and radiochemical purity in preliminary studies were (>68%, >99%, >95%), respectively.

### PET/CT, MR imaging and biodistribution

For PET/CT imaging, mice were anesthetized using 3.0% isoflurane in oxygen, maintained at 1.5–2.0% isoflurane, and subjected to ^64^Cu-C4BP injections (126 ± 22 mCi (4.6±0.8 MBq)/mouse). Images of *KT;Ptf1a^CreER^;Trp53^fl/fl^* mice (n=6) and age-matched wild type C57BL/6 mice (n=6) were acquired at 0, 4, and 21 hours post-injection. PET/CT (Inveon, Siemens) scanning for the 0-hour time point started on the PET scanner bed with a simultaneous tail-vein injection of ^64^Cu-C4BP for 30 minutes to obtain dynamic temporal images, then subsequently, CT imaging was acquired for 10 minutes for the anatomical reference. The following time-point scans were performed at 4 and 21 hours. All 12 mice were scanned at 49 and 84 days after tamoxifen treatment to induce tumors. At 84 days, 6 mice (n=3: *KT;Ptf1a^CreER^;Trp53^fl/fl^* mice, n=3 wild-type control mice) were euthanized after PET/CT scan, and *ex vivo* PET imaging of organs such as liver, spleen, pancreas, colon and small intestines was performed. After collecting all organs of interest (blood, heart, lungs, spleen, stomach feces, kidneys, muscle, bone, skin, and brain), radioactivity in each organ was measured using a gamma counter. The biodistribution of ^64^Cu-C4BP is presented as percent injected activity per g (%IA/g) in organs. At 120 days, the remaining mice were imaged with PET/CT, and based on morbidity, were euthanized. One mouse showed significant weight loss was scanned under PET/CT and T2-weighted), and bright field image of abdominal area was taken. T2-weighted MR images were acquired with an 11.7 T Agilent MR scanner with Bruker interface. Mice were oriented similar to PET/CT and scanned with a TR of 2500 msec and a TE of 3 msec with standard fat suppression T2-weighted fast spin-echo sequence with a slice thickness of 0.8 mm to capture the pancreas. Organs showing peritoneal metastasis were further analyzed with IHC and H&E.

### Quantitative region of interest (ROI) analysis

PET image analysis was performed with Inveon Research Workplace (IRW) software after the co-registration of PET and CT images. Images were quantified by manually drawing regions of interest (ROI) in the hot spots adjacent to abdominal organs such as stomach, liver and kidney. Radioactivity in organs was normalized to the percent injected activity per cubic centimeter volume (%IA/cc).

### Software

Rstudio with R version 4.2.2 was used for computation environment and language. Seurat v4 was used for spatial transcriptomic analysis and clusterProfiler v4 was used for enrichment analysis. Monocle3 was used for pseudotime analysis. Ggplot was used for plotting analysis results in R. Custom scripts were created for data wrangling between packages, MA plots, surfaceome score calculations, and mapping of pseudotime onto spatial dimension. Enable Medicine’s online proprietary software and R package, Spatialmap, were used for multiplexed IHC analysis.

### Statistical analyses

Statistical analyses were completed using Graphpad Prism 9 (RRID:SCR_000306) and RStudio 2023.09.1 and R version 4.2.2. Significance was assessed using an unpaired t-test and one-way ANOVA as described in the figure captions. Pearson’s correlations were calculated with the correlation function in R (corr).

### Data availability

Data are available upon reasonable request. Source data will be provided with this paper.

## Acknowledgements

The authors acknowledge the support of NIGMS5T32GM007276, NIHR01CA250557, and R01CA253316. The assistance of the Attardi and Artandi laboratories in the mouse model generation and interpretation is greatly appreciated. We thank Dr. Jane Chen at UC Davis Center for Genomic Pathology Lab for mouse tissue process of IHC and H&E. We also thank Peter Chou, Aaron Chiou, and Aya Kondo for guidance using Enable Medicine’s cloud platform. PET/CT/MRI imaging was performed at the Stanford Center for Innovation in In Vivo Imaging (SCi3) and Canary Center of Stanford University.

## Author contributions

J.W. A.J.K., J.W.S. and K.W.F. designed and implemented the study and wrote the paper. G.C. and M.S. provided samples and pathological interpretation. W.P., R.B. and A.I. provided clinical guidance on the study design and image interpretation. M.P., S.T., B.W. and M.R. performed experiments.

## Competing interests

The authors declare no competing interests.

## Notes

### Competing Interest Statement

The authors have declared no competing interest.

