## Supplement for "Spatial transcriptomic analysis drives PET imaging of tight junction protein expression in pancreatic cancer theranostics"

### Supplementary Methods

### Supplementary Figures

### Supplementary Tables

**Supplementary Methods**

**Xenograft mouse model**: Xenograft tumors were established in both hind flanks of NOD/SCID mice (Jackson Laboratory) by subcutaneous injection of Capan-1 cells ($1 \times 10^6$) suspended in 1:1 Matrigel:DMEM medium (100 μL).

**PET/CT imaging and ROI analysis of xenograft mice:** The PET/CT imaging of xenograft NOD/SCID (Jackson Laboratory) mice (n=4) bearing Capan-1 tumors followed the scanning protocol in the Online Methods Section once tumor size reached ~5 mm diameter. Mice (n=4) received a tail vein injection of $^{64}$Cu-C4BP (94±11μCi or 3.5±0.4 MBq/mouse). For blocking (n=4), injection of $^{64}$Cu-C4BP was proceeded with ~1000-fold excess cold compound (100 μg). Accumulation of radioactivity (tumor, kidneys, and blood) were quantified by manually drawing regions of interest (ROI) after co-registration of PET/CT images on Inveon Research Workplace (IRW) software. Organ radioactivity was normalized to the percent injected activity per cubic centimeter volume (%IA/cc).

**Western blot of CLDN4:** The tumor cell lines Capan-1, PANC-1, mT4, and NDL were lysed in RIPA lysis buffer (ThermoFisher Scientific) for 10 minutes at 4 °C. After protein quantification via the bicinchoninic acid (BCA) assay, protein concentrations were standardized to 0.4 mg/mL. The homogenized protein samples were then combined with the loading buffers (ThermoFisher Scientific) per the manufacturer's guidelines, and 2 μg of each sample was loaded per well on an SDS-PAGE gel. Following SDS-PAGE separation, proteins were transferred to a PVDF membrane (ThermoFisher Scientific). For protein detection, the membrane was stained with primary antibodies: a rabbit polyclonal antibody against CLDN4 (1:200 dilution, ThermoFisher Scientific, Catalog #36-4800) and a rabbit antibody against β-actin (1:2000 dilution, Sigma Aldrich). Primary antibody bound to each protein was stained by incubation with a secondary goat anti-rabbit IgG (H+L) antibody conjugated with horseradish peroxidase (HRP) (1:2000 dilution, Thermo Fisher Scientific). Visualization of protein bands was done using a chemiluminescence substrate (BioRed) and imaging system (BioRad, ChemiDoc XRS+ CCD) for subsequent analyses.

**Immunohistochemistry of CLDN4***:* Harvested tissues were fixed with 10% formalin solution for more than 24 hours and then processed using the standard paraffin embedded tissue sectioning procedure by the UC Davis Center for Genomic Pathology Lab. Briefly, antigen retrieval was performed for 45 min with a citrate buffer at pH 6.0 in a Decloaking Chamber (Biocare Medical, Concord, CA) at 125°C and 15 psi. Slides were blocked with normal goat serum then incubated with a hCLDN4 polyclonal rabbit antibody (1:200, ThermoFisher Scientific, Catalog #36-4800) overnight at room temperature in a humidified chamber, followed by a biotinylated goat anti-rabbit secondary antibody (1:1000, Vector Labs, BA-1000-1.5). The Vectastain ABC Kit Elite Kit and a diaminobenzidine Peroxidase Substrate Kit (Vector Labs, Burlingame, CA) were used for amplification and visualization of signal, respectively.

**H&E**: Tissues were fixed in 10% neutral buffered formalin at room temperature for 24 h then placed in 70% ethanol until processing, which was normally within 24 h. A Tissue-Tek VIP autoprocessor (Sakura, Torrance, CA) was used to process tissues that were then embedded in Paraplast (melting temperature 56–60°C), sectioned to 4 μm and mounted on glass slides. Sections were stained with hematoxylin and eosin (H&E) for pathologic analysis.

**KEGG Pathway analysis**

KEGG pathway enrichment analysis was performed with a custom wrapper function that combined Seurat findMarker function to extract differential expressions and the pathview function to map differential expression fold change values onto native KEGG maps. Key components in the native KEGG maps were selected to plot the KEGG derived pathways with fold change values mapped onto each node.

### Supplementary Figures

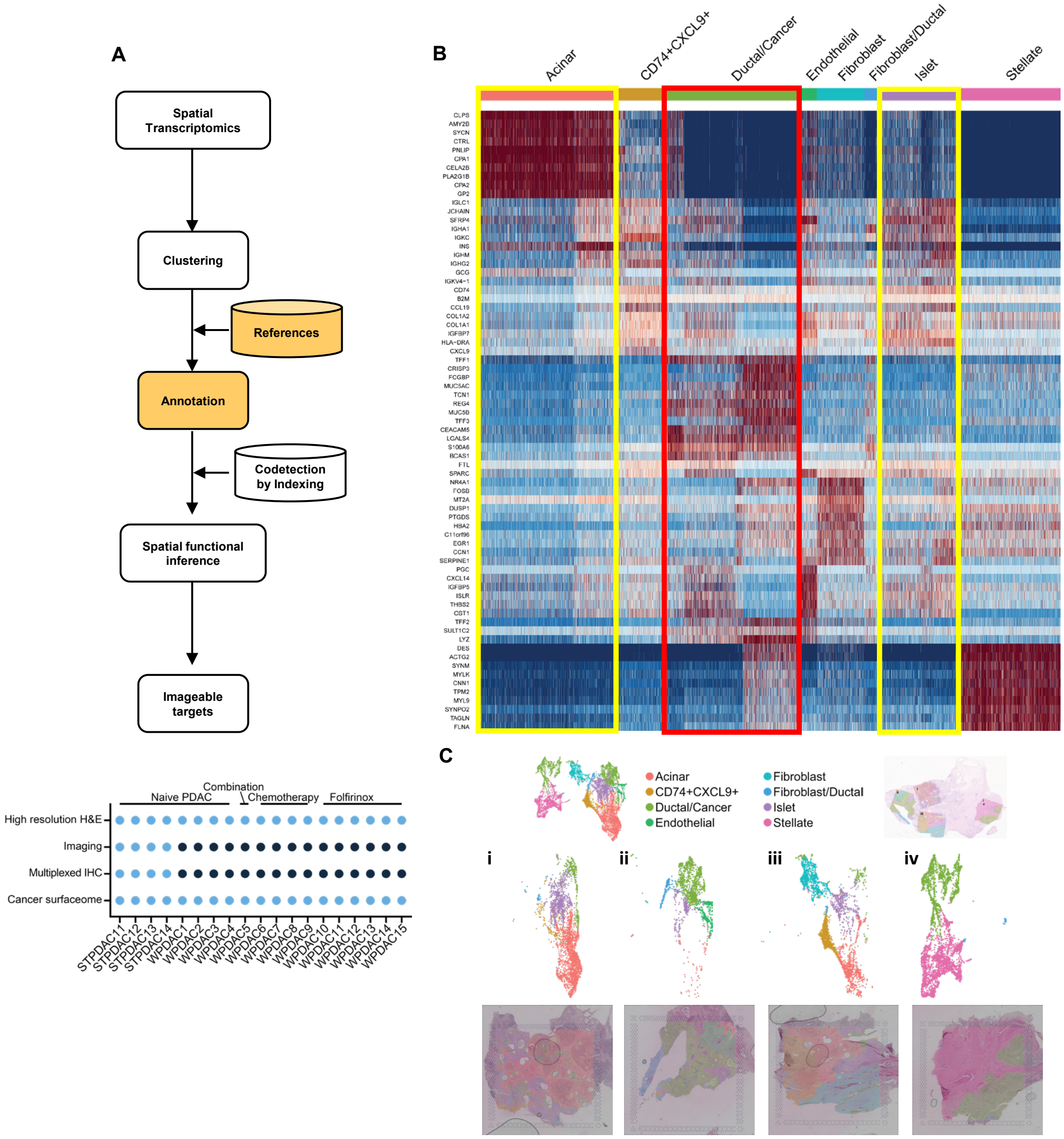


**Supplementary Fig. S1. Heatmap for all pancreatic clusters in PDAC and the corresponding cell types.** A) Top) Summary method schematic of the paper. Bottom) Summary of the PDAC samples and corresponding treatments. B) Heatmap of top cluster top differentially expressed genes and annotated based on scRNA sequencing. C) Clustering and overlap of clusters on histology sections.

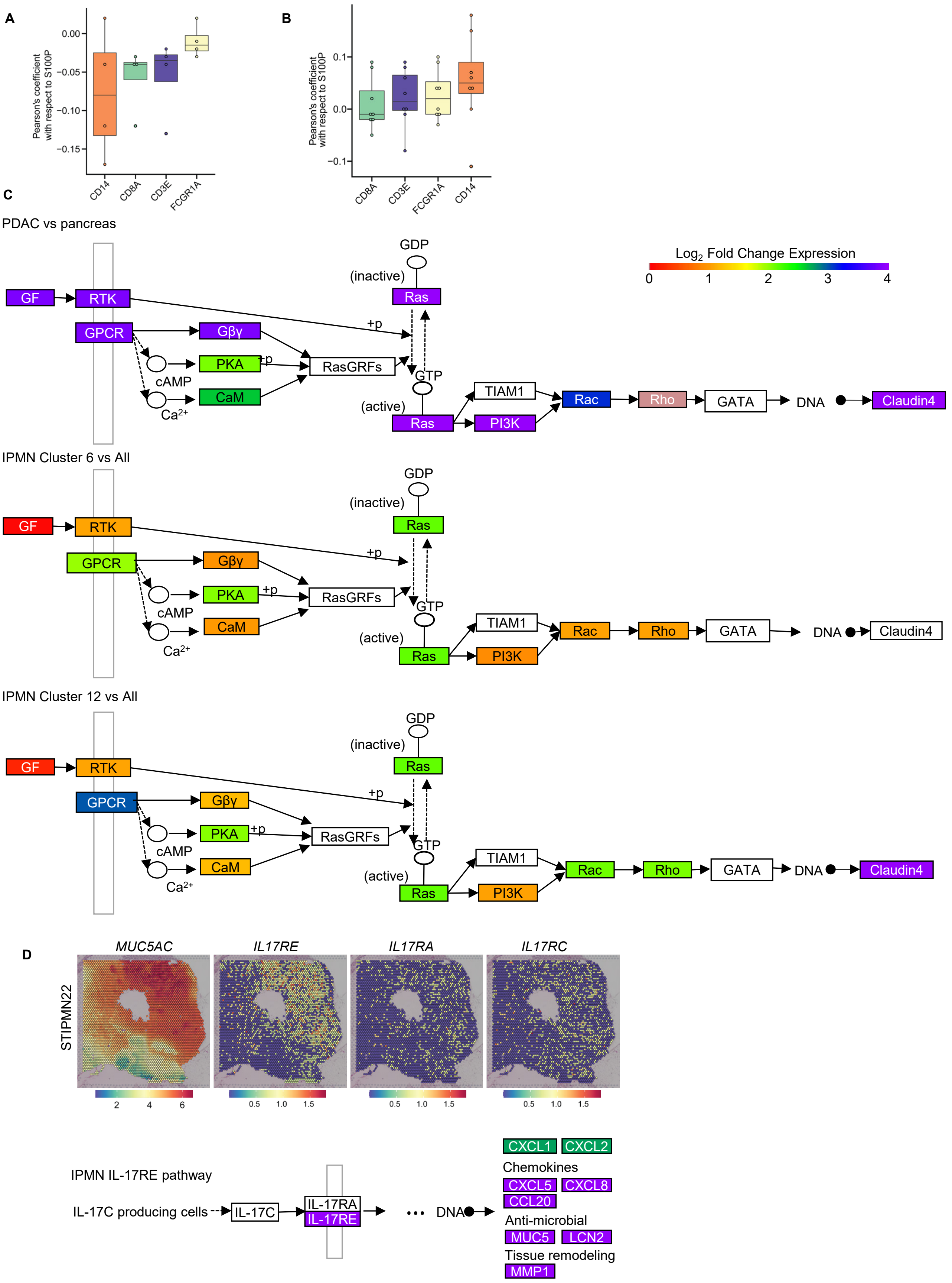


**Supplementary Fig. S2. Correlation of immune cell markers with S100P and underlying pathways for *Claudin-4* and *MUC5AC* expression.** A-B) Pearson's coefficient for the spatial correlation of immune cell markers (*CD3, CD8, CD64 (FCGR1A), CD14*) and S100P. C) Pathways derived from KEGG associated with claudin-4, D) IL-14RE pathway and MUC5AC.

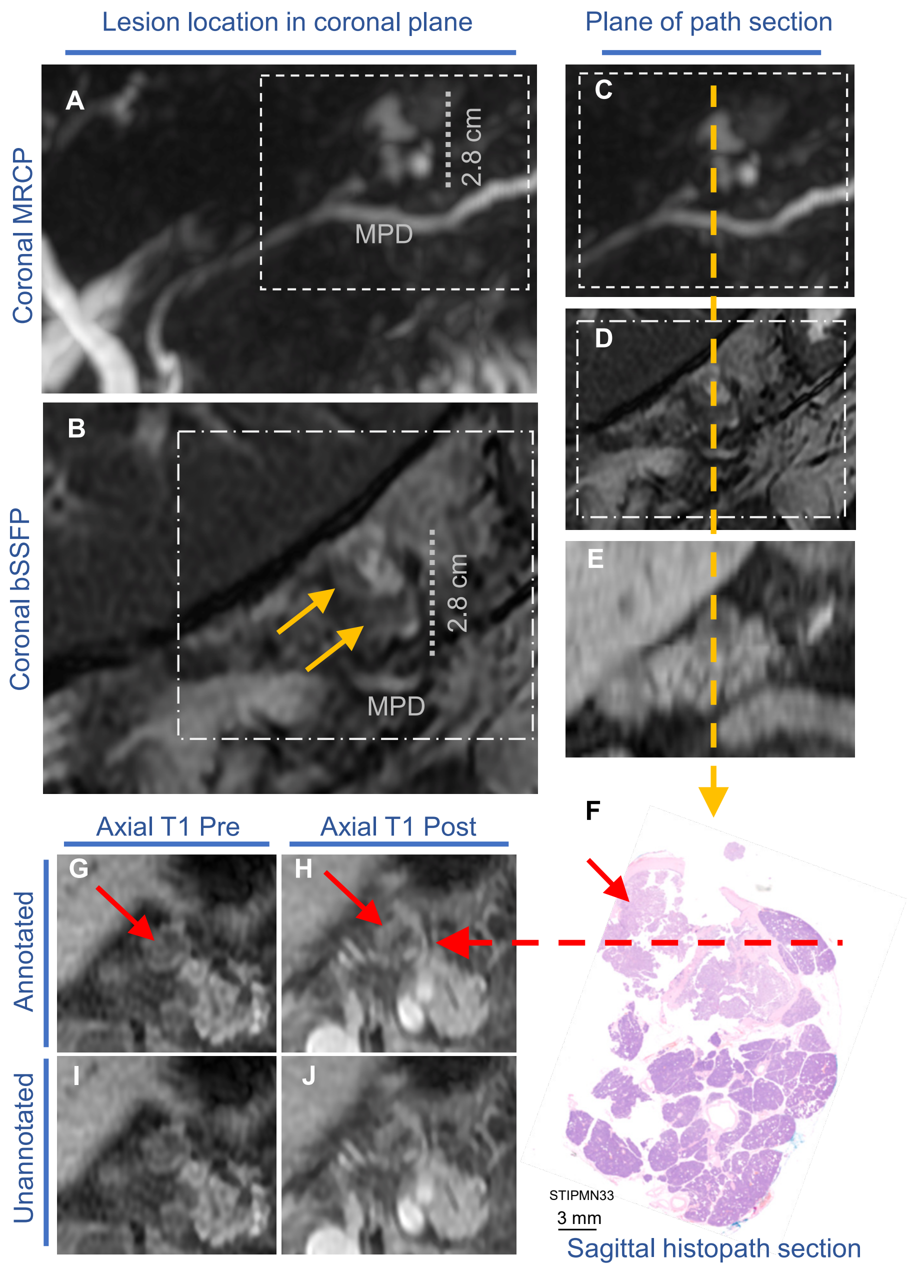


**Supplementary Fig. S3. Corresponding magnetic resonance imaging (MRI) of IPMN sections near STIPMN34 tissue.** A) Coronal balanced steady state free precession (bSSFP) and B) magnetic resonance cholangiopancreatography (MRCP) MRI axial T1-weighted slices of the pancreas near the STIPMN34 location (lesion is annotated with yellow arrows). C-E) Enhanced-resolution images corresponding with the plane of the pathology section (F) and referenced to (A) and (B) by the image border. G-J) In axial T1-weighted imaging pre and post contrast injection, enhancement of the lesion corresponds with the region demonstrating an elevated cancer surfaceome score and elevated CLDN4 (red arrows are corresponding locations).

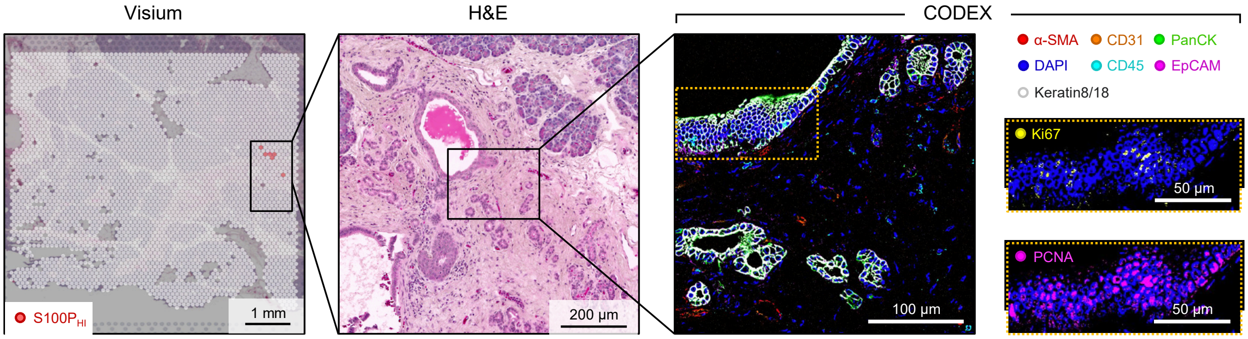


**Supplementary Fig. S4. Spatial transcriptomics/CODEX imaging of small ductal lobules demonstrating sensitivity of each technique.** Combining spatial proteomics and transcriptomics enhanced the detection of small ductal lobules that were located in the pancreas surrounding IPMN lesions. These small lesions were considered to be precancers based on nuclear morphology and H&E. In spatial transcriptomics, these lesions expressed *S100P* and *MUC5AC* but not *CLDN4*.

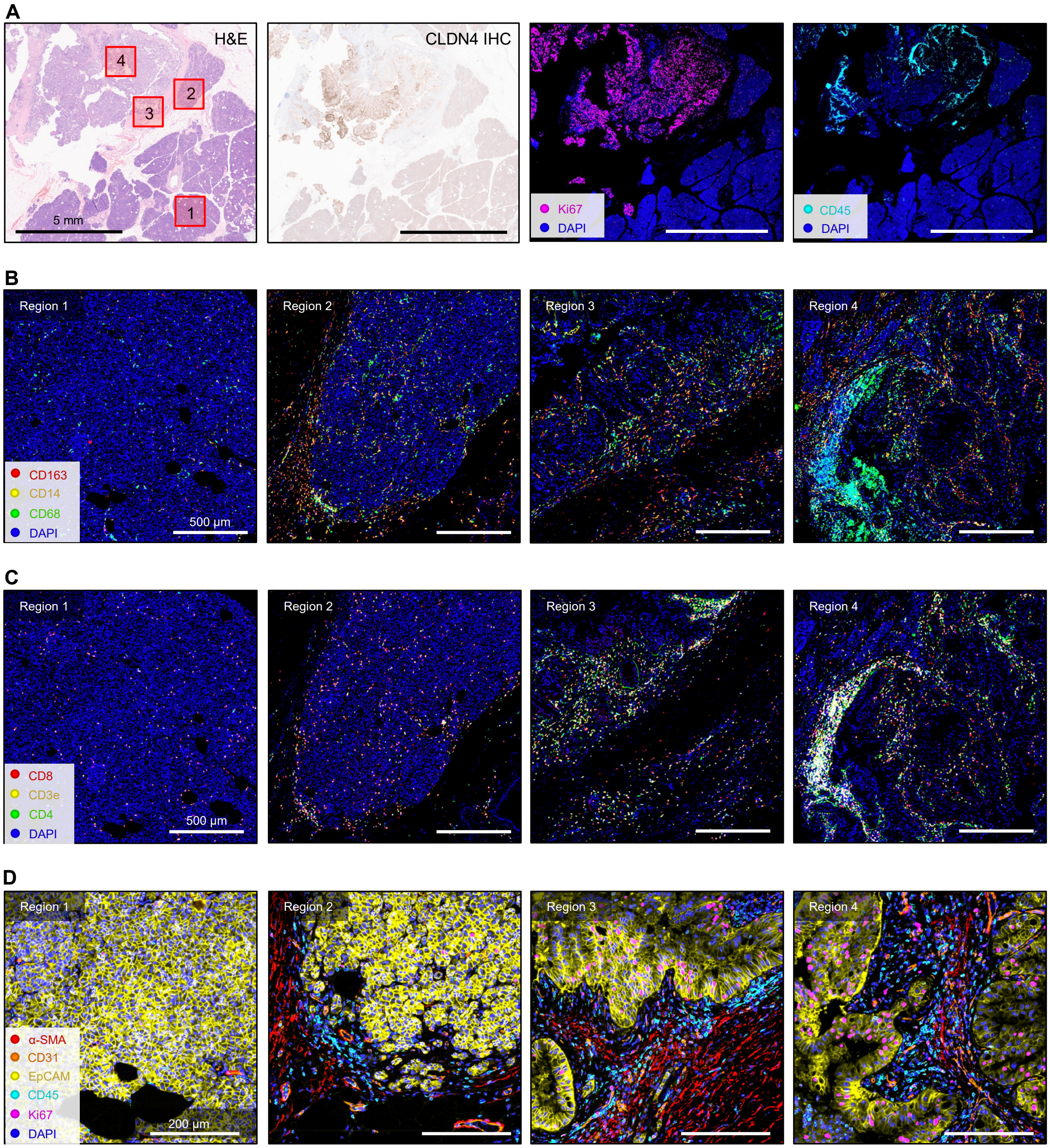


**Supplementary Fig. S5. CODEX imaging of IPMN.** A) Region layout of H&E and CLDN4 IHC, with Ki67 and CD45 CODEX expression maps. B-C) Regional mapping showing B) myeloid markers and C) T cell markers. D) Regional zoom in showing epithelial, proliferation, and vessel marker expression, demonstrating regional variability throughout the IPMN.

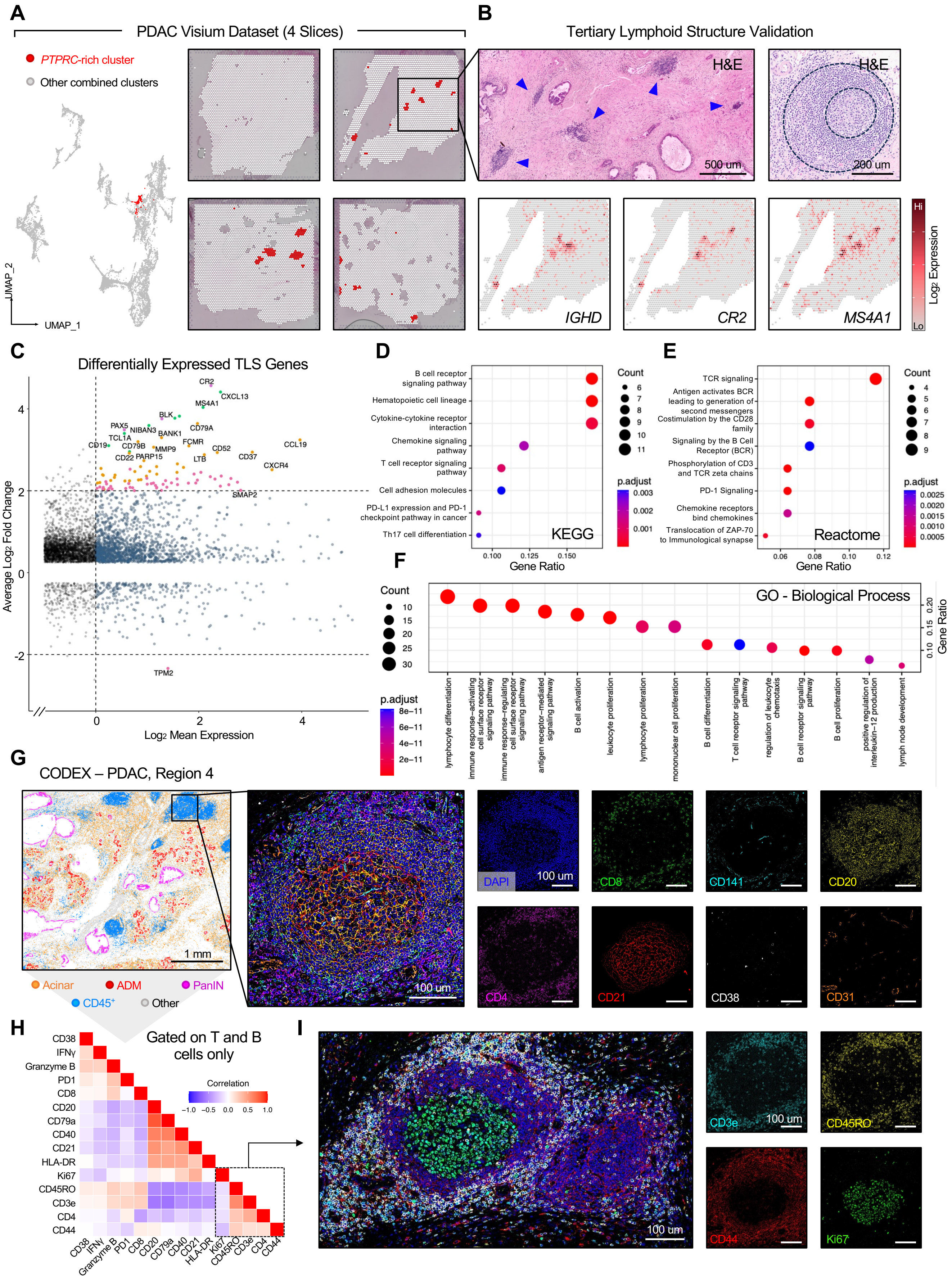
A
PDAC Visium Dataset (4 Slices)
PTPRC-rich cluster
Other combined clusters
UMAP_2
UMAP_1
B
Tertiary Lymphoid Structure Validation
H&E
500 um
H&E
200 um
IGHD
CR2
MS4A1
Hi
Lo
Log2 Expression
C
Differentially Expressed TLS Genes
CR2
CXCL13
MS4A1
BLK
PAX5
NIBAN3
CD79A
TCL1A
BANK1
CD19
CD79B
FCMR
MMP9
CD52
CCL19
CD22
PARP15
LTB
CD37
CXCR4
SMAP2
TPM2
Average Log2 Fold Change
Log2 Mean Expression
D
B cell receptor signaling pathway
Hematopoietic cell lineage
Cytokine-cytokine receptor interaction
Chemokine signaling pathway
T cell receptor signaling pathway
Cell adhesion molecules
PD-L1 expression and PD-1 checkpoint pathway in cancer
Th17 cell differentiation
KEGG
Gene Ratio
Count
p.adjust
E
TCR signaling
Antigen activates BCR leading to generation of second messengers
Costimulation by the CD28 family
Signaling by the B Cell Receptor (BCR)
Phosphorylation of CD3 and TCR zeta chains
PD-1 Signaling
Chemokine receptors bind chemokines
Translocation of ZAP-70 to Immunological synapse
Reactome
Gene Ratio
Count
p.adjust
F
GO - Biological Process
Count
p.adjust
Gene Ratio
lymphocyte differentiation
immune response−activating cell surface receptor signaling pathway
immune response−regulating cell surface receptor signaling pathway
antigen receptor−mediated signaling pathway
B cell activation
leukocyte proliferation
lymphocyte proliferation
mononuclear cell proliferation
B cell differentiation
T cell receptor signaling pathway
regulation of leukocyte chemotaxis
B cell receptor signaling pathway
B cell proliferation
positive regulation of interleukin−12 production
lymph node development
G
CODEX – PDAC, Region 4
1 mm
100 um
Acinar
ADM
PanIN
CD45+
Other
DAPI
100 um
CD8
CD141
CD20
CD4
CD21
CD38
CD31
H
Gated on T and B cells only
Correlation
CD38
IFNγ
Granzyme B
PD1
CD8
CD20
CD79a
CD40
CD21
HLA-DR
Ki67
CD45RO
CD3e
CD4
CD44
I
100 um
CD3e
100 um
CD45RO
CD44
Ki67

**Supplementary Fig. S6. CODEX imaging of tertiary lymphoid organs (TLOs).** A) Four spatial transcriptomic PDAC regions were merged, normalized with the SCTransform function, and clustered with Seurat. The *protein tyrosine phosphatase receptor type C* (*PTPRC, CD45*)-rich cluster in the dataset was highlighted in red both on Uniform Manifold Approximation and Projection (UMAP) and in spatial transcriptomic coordinates, corresponding to TLOs. B) Overlapping H&E and B-cell gene signatures confirm the presence of TLOs (blue arrowheads). C) Magnitude and average (MA) plot of differentially-expressed genes (DEGs) in the *PTPRC*-rich/TLO cluster in comparison to the rest of the PDAC spatial transcriptomic dataset. D) KEGG, E) REACTOME, and F) GO – Biological Process Pathway analysis of DEGs > average $\log_2$ fold change of 2 from C). G) CODEX imaging of TLO and several immune surface receptors from a representative PDAC region. H) Pearson's correlation of several surface markers expressed on T and B cells from the PDAC region in G). I) CODEX imaging of the memory T-cell shell and proliferating B-cell core of a TLO, validating the Pearson correlation from H).

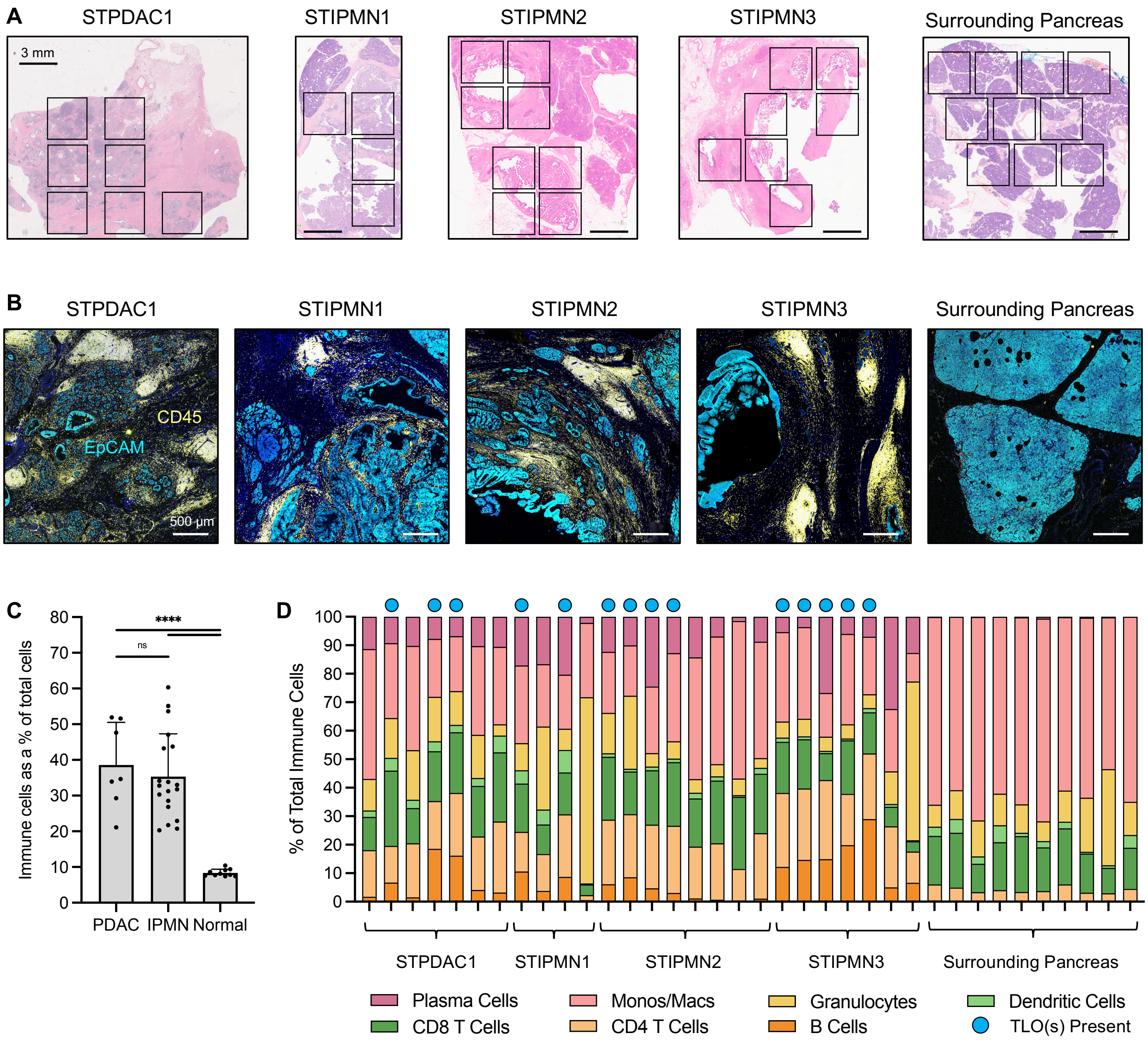


**Supplementary Fig. S7. CODEX imaging and analysis of immune cells across PDAC and IPMN.** A) IHC of target regions. B) CODEX imaging of DAPI (dark blue), EpCAM (light blue) and CD45 (yellow) for regions of PDAC, IPMN and surrounding pancreas. C) Overall fraction of immune cells across all fields. D) Quantification of immune cell phenotype for varied fields and identification of regions containing at least 1 TLO. One-way ANOVA results are as follows for C): **** = p<0.0001, ns = not significant.

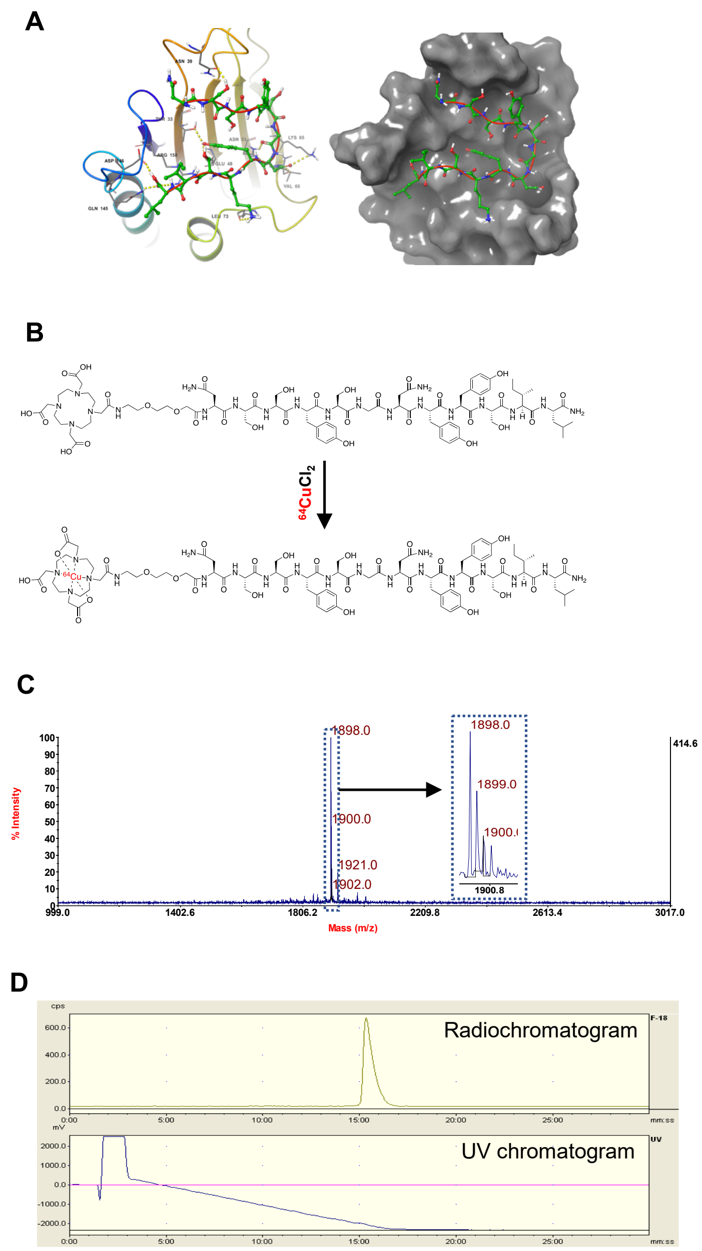


**Supplementary Fig. S8. Peptide targeting of claudin-4.** A) Modeling demonstrates the NSSYSGNYYSIL peptide ($K_D$, 1.97 nM) in binding pocket. B) $^{64}$Cu-labeling scheme. C) MALDI spectrum of DOTA-PEG$_1$-C4BP. D) Radio- and UV-chromatogram of $^{64}$Cu-C4BP.

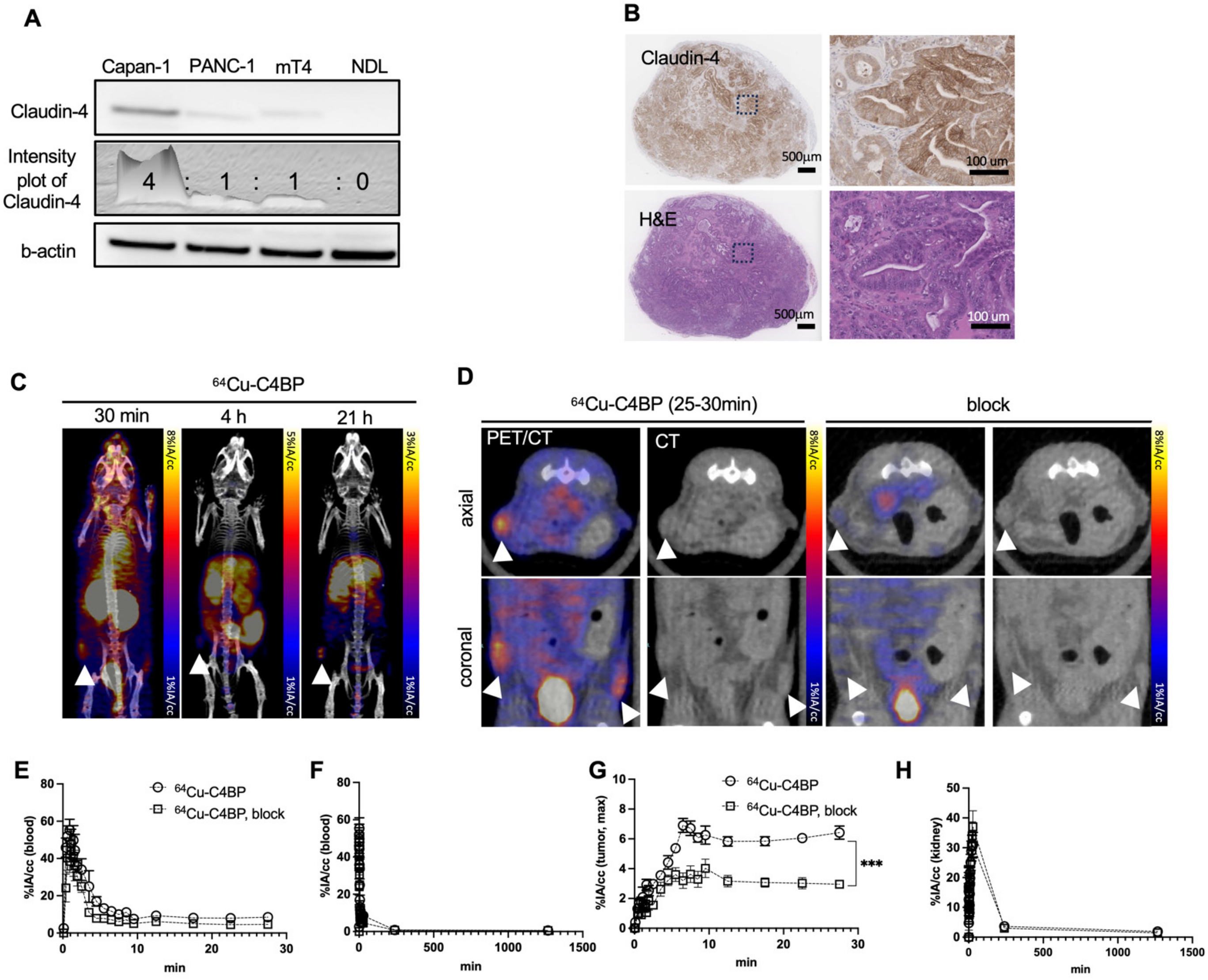


**Supplementary Fig. S9. Molecular imaging of claudin-4 in a human xenograft mouse model.** A) Expression of claudin-4 in various cell lines (Capan-1: human pancreatic adenocarcinoma, PANC-1: human pancreatic cancer cell line, mT4: murine pancreatic cancer cell line, NDL: mouse mammary carcinoma cell line). B) CLDN4 IHC and H&E in Capan-1 tumors. C) Representative MIP PET/CT image of xenograft mouse bearing Capan-1 tumors. Image represents $^{64}$Cu-C4BP retention in tumors over time. D) PET/CT slice images with and without co-injection of 1000-fold excess of cold peptide for blocking. E-H) Time activity curves (TACs) for $^{64}$Cu-C4BP concentration in E-F) blood, G) tumor and H) kidney. Abbreviations: B: bladder, K: kidney, H: heart, L: liver. * = $p<0.05$ in G). Unpaired two-tailed t-test was performed in G (*** = $p<0.05$).

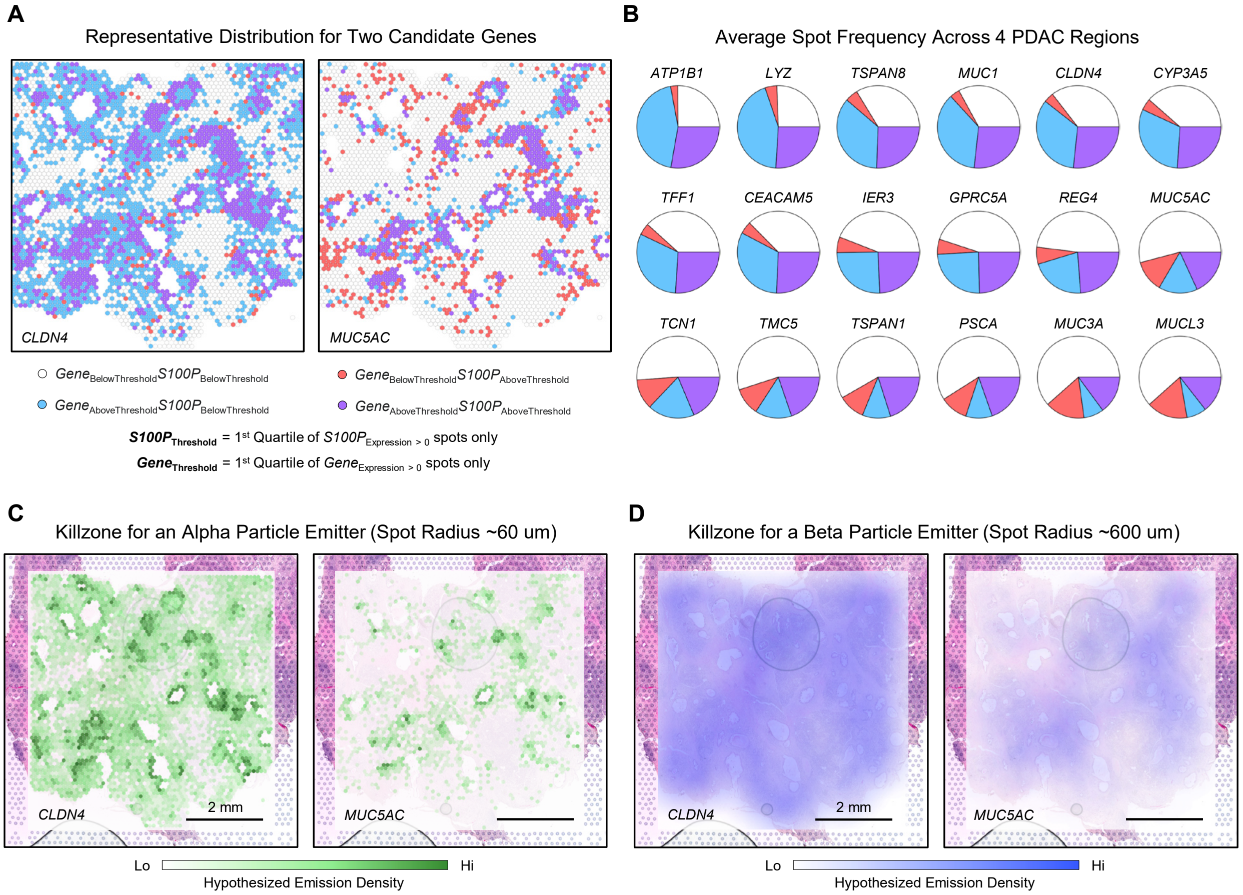


**Supplementary Fig. S10. Modeling of claudin-4 radiopharmaceutical distribution for future therapeutic applications.** A) Visium spots for a representative PDAC region were colored using the 1st quartile of gene/*S100P* expression as a threshold, demonstrating that a claudin-4/mucin-5ac specific theragnostic agent would detect all S100P expressing cells (purple) and additional cells (blue). The spots in red contain *S100P* but not *CLDN4* or *MUC5AC*, representing areas that would be missed after administering a theragnostic. B) Similar analysis to A) for select cancer-upregulated genes averaged across four PDAC regions. C-D) Kill region for CLDN4 and MUC5AC alpha (C) and beta (D) emitters, assuming all regions were accessible to the peptide.

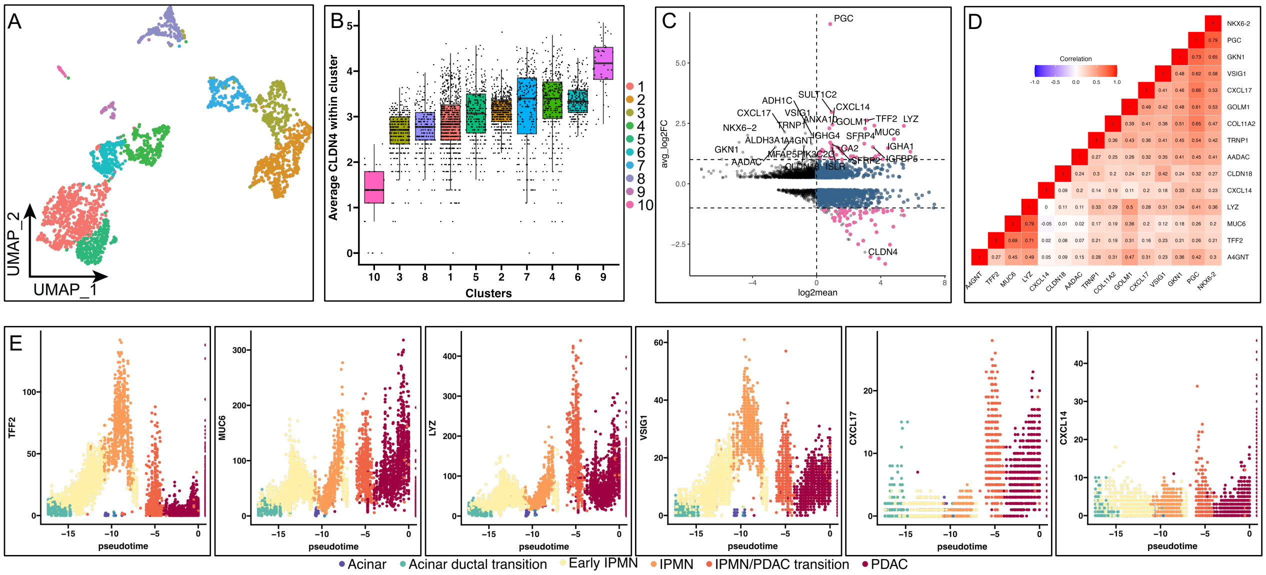


**Supplementary Fig. S11. Analysis of cellular subsets within Cluster 1 (Fig. 1).** A) UMAP of subclusters. B) CLDN4 expression within subclusters. C) Differential expression for subcluster 10 as compared with the remainder of cluster 1. D) Pearson's correlation of mRNA expression within genes enhanced in subcluster 10 (lower CLDN4 expression). E) Pseudotime analysis of genes with enhanced expression in subcluster 10 (*TFF2*, *MUC6*, *LYZ*, *VSIG1*, *CXCL17*, *CXCL14*).

#### Supplementary Tables

**Supplementary Table S1.** Clinical data regarding patient samples used in this study acquired at Stanford University and from Ref. 20 and 21.

| Sample | Gender | Age | Grade | Location | Diabetes |
|---|---|---|---|---|---|
| LG_1 | F | 75 | LG IPMN | head | No |
| LG_2 | M | 76 | LG IPMN | head | Yes |
| LG_3 | M | 56 | LG IPMN | head, body | Yes |
| LG_4 | F | 67 | LG IPMN | head | Yes |
| LG_5 | F | 76 | LG IPMN | body/tail | Yes |
| LG_6 | F | 51 | LG IPMN | body/tail | No |
| LG_7 | M | 76 | LG IPMN | body/tail | No |
| HG_1 | M | 72 | HG IPMN | head | No |
| HG_2 | F | 89 | HG IPMN | head | No |
| HG_3 | M | 62 | HG IPMN | head | No |
| PDAC_1 | F | 76 | PDAC | Head | No |
| PDAC_2 | M | 85 | PDAC | Head, Uncinate | Yes |
| PDAC_3 | F | 66 | PDAC | head | Ues |
| HT264P1 | F | 77 | PDAC | body | No |
| HT270P1 | M | 60 | PDAC | Head | Type II |
| HT284P1 | M | 76 | PDAC | neck/body margin | Type II |
| HT288P1 | M | 73 | PDAC | Head | Type II |
| HT224P1 | F | 61 | PDAC | Head | Type II |
| HT231P1 | F | 63 | PDAC | Tail | No |
| HT231P1-2 | F | 63 | PDAC | Tail | No |
| HT232P1 | F | 60 | PDAC | body/tail | No |
| HT232P1-2 | F | 60 | PDAC | body/tail | No |
| HT259P1 | F | 69 | PDAC | Head | No |
| HT306P1 | F | 74 | PDAC | Head | No |
| HT306P1-2 | F | 74 | PDAC | Head | No |
| HT306P1-3 | F | 74 | PDAC | Head | No |
| HT306P1-4 | F | 74 | PDAC | Head | No |
| HT242P1 | M | 70 | PDAC | Tail (proximal remanant) | Yes |
| STPDAC11 | M | 56 | PDAC | Head | Type II (2 mos) |
| STPDAC12 | M | 56 | PDAC | Head | Type II (2 mos) |
| STPDAC13 | M | 56 | PDAC | Head | Type II (2 mos) |
| STPDAC14 | M | 56 | PDAC | Head | Type II (2 mos) |
| STIPMN11 | F | 82 | HG IPMN | Head | Prediabetes |
| STIPMN12 | F | 82 | HG IPMN | Head | Prediabetes |
| STIPMN21 | M | 63 | HG IPMN | Body/tail | Type II (2 mos) |
| STIPMN22 | M | 63 | HG IPMN | Body/tail | Type II (2 mos) |
| STIPMN31 | M | 73 | HG IPMN | Tail | No |
| STIPMN32 | M | 73 | HG IPMN | Tail | No |
| STIPMN33 | M | 73 | HG IPMN | Tail | Prediabetes |
| STIPMN34 | M | 73 | HG IPMN | Tail | Prediabetes |

**Supplementary Table S2.** Genes from cancer surfaceome with greater than 80% frequency of expression in pancreatic cancer.

| | |
|---|---|
| B2M | 100.00 |
| EEF1A1 | 99.99 |
| S100A6 | 99.94 |
| HLA-A | 99.35 |
| CD63 | 98.65 |
| ITM2B | 98.63 |
| HLA-C | 98.41 |
| IFITM3 | 98.34 |
| SPINT2 | 97.90 |
| RAC1 | 97.73 |
| HSP90AA1 | 97.55 |
| CD74 | 97.29 |
| EPCAM | 96.18 |
| TM4SF1 | 96.16 |
| CYBA | 96.10 |
| CD59 | 95.89 |
| RPSA | 95.73 |
| HSP90AB1 | 94.15 |
| ATP1A1 | 93.57 |
| ATP1B1 | 93.26 |
| VAMP8 | 92.40 |
| LY6E | 92.10 |
| TSPAN8 | 90.40 |
| ARF1 | 89.77 |
| HLA-E | 89.27 |
| RHOA | 89.18 |
| CDC42 | 89.04 |
| CLDN4 | 88.83 |
| CD151 | 88.61 |
| TACSTD2 | 88.23 |
| CIB1 | 87.51 |
| IFITM2 | 87.32 |
| CD164 | 86.53 |
| ITGB1 | 86.47 |
| APLP2 | 86.33 |
| CD9 | 86.33 |
| RTN4 | 86.27 |
| ENO1 | 86.15 |
| UBE2D3 | 86.12 |
| CALR | 85.88 |
| RAB11A | 85.57 |
| APP | 85.51 |
| MUC1 | 85.32 |
| TSPAN3 | 85.24 |
| EZR | 85.06 |
| BSG | 85.03 |
| VDAC1 | 84.98 |
| P4HB | 84.87 |
| HLA-DRA | 84.85 |
| TMEM123 | 84.52 |
| TMED10 | 83.19 |
| VAPA | 81.93 |
| VPS28 | 81.55 |
| TMEM219 | 81.32 |
| MAL2 | 81.02 |
| CLDN7 | 80.28 |
| HLA-B | 80.05 |

**Supplementary Table S3**. Biodistribution of $^{64}$Cu-C4BP (given as %IA/g) in organs of interest at 22h.

| | KT;Ptf1a$^{CreER}$;Trp53$^{fl/fl}$ (n=3) | | Wild type (n=3) | | |
|---|---|---|---|---|---|
| | Mean | Stdev | Mean | Stdev | P value |
| blood | 1.60 | 1.84 | 0.20 | 0.01 | 0.256 |
| heart | 1.23 | 0.59 | 0.29 | 0.05 | 0.051 |
| lungs | 0.89 | 0.09 | 0.50 | 0.20 | 0.036 |
| liver | 7.73 | 5.35 | 1.80 | 0.56 | 0.129 |
| pancreas | 1.46 | 0.86 | 0.16 | 0.02 | 0.058 |
| spleen | 1.27 | 0.63 | 0.24 | 0.03 | 0.045 |
| stomach | 0.54 | 0.24 | 0.60 | 0.16 | 0.778 |
| small intestine | 2.45 | 1.31 | 0.80 | 0.15 | 0.095 |
| large intestine | 4.08 | 2.64 | 0.90 | 0.17 | 0.106 |
| feces | 1.92 | 1.12 | 1.13 | 0.49 | 0.324 |
| kidneys | 6.57 | 3.85 | 1.14 | 0.13 | 0.071 |
| muscle | 0.38 | 0.16 | 0.10 | 0.01 | 0.041 |
| bone (femur) | 0.55 | 0.25 | 0.13 | 0.08 | 0.051 |
| skin | 0.78 | 0.43 | 0.20 | 0.04 | 0.080 |
| brain | 0.16 | 0.08 | 0.06 | 0.00 | 0.088 |
| pancreas/muscle | 3.71 | 0.56 | 1.53 | 0.10 | 0.003 |

**Supplementary Table S4**. Genes with enhanced expression in subcluster 10 compared with the remainder of cluster 1 from Figure 1 (restricted to genes with average $log_2$ fold change greater than 1.5 and adjusted p value below 0.05).

| *Gene* | Average $log_2$ fold change | Adjusted p value |
|---|---|---|
| *PGC* | 6.61080985 | 6.57E-34 |
| *SULT1C2* | 2.9686981 | 1.30E-16 |
| *TFF2* | 2.62198568 | 3.28E-14 |
| *VSIG1* | 2.55354617 | 8.11E-23 |
| *CXCL14* | 2.44793598 | 2.43E-13 |
| *MUC6* | 2.40624072 | 1.85E-13 |
| *LYZ* | 2.3957633 | 1.11E-14 |
| *ANXA10* | 2.29393608 | 5.76E-16 |
| *GOLM1* | 2.28199427 | 1.29E-15 |
| *CXCL17* | 2.13658978 | 9.47E-36 |
| *TRNP1* | 2.08757002 | 1.26E-19 |
| *ADH1C* | 2.02782329 | 4.67E-18 |
| *NKX6-2* | 2.010595 | 2.25E-226 |
| *A4GNT* | 1.8987742 | 3.17E-29 |
| *IGHA1* | 1.84919811 | 2.05E-10 |
| *GKN1* | 1.71511309 | 0 |
| *MFAP5* | 1.71138147 | 1.21E-22 |
| *ALDH3A1* | 1.7084192 | 3.05E-27 |
| *IGHG4* | 1.70636113 | 9.34E-11 |
| *SFRP4* | 1.65802546 | 3.89E-10 |
| *PIK3C2G* | 1.64872981 | 2.08E-14 |
| *CA2* | 1.6393763 | 1.80E-13 |
| *CLDN18* | 1.56122327 | 2.60E-15 |
| *ISLR* | 1.54309321 | 4.99E-08 |
| *AADAC* | 1.51937112 | 1.57E-41 |
| *IGFBP5* | 1.5173545 | 1.66E-06 |
| *SFRP2* | 1.51135135 | 1.01E-07 |
